# Ancient DNA Reveals China as a Historical Genetic Melting Pot in Tiger Evolution

**DOI:** 10.1101/2022.09.14.507899

**Authors:** Xin Sun, Yue-Chen Liu, Mikhail P. Tiunov, Dmitry O. Gimranov, Yan Zhuang, Yu Han, Carlos A. Driscoll, Yu-Hong Pang, Chunmei Li, Yan Pan, Marcela Sandoval Velasco, Shyam Gopalakrishnan, Rui-Zheng Yang, Bao-Guo Li, Kun Jin, Xiao Xu, Olga Uphyrkina, Yan-Yi Huang, Xiao-Hong Wu, M. Thomas P. Gilbert, Stephen J. O’Brien, Nobuyuki Yamaguchi, Shu-Jin Luo

## Abstract

The contrast between the tiger’s (*Panthera tigris*) 2-3 My age and extant tigers’ coalescence approximately 110,000 years ago suggests an ancient demographic bottleneck. Here we collected over 60 extinct specimens across mainland Asia and generated whole genome sequences from a 10,600-year-old Russian Far East (RFE) specimen (RUSA21, 8× coverage), 14 South China tigers (0.1-12×), three Caspian tigers (4-8×), plus 17 new mitogenomes. RUSA21 clustered within modern Northeast Asian phylogroups and partially derived from an extinct Late Pleistocene lineage. While some 8,000-10,000-year-old RFE mitogenomes are basal to all tigers, one 2,000-year-old specimen resembles present Amur tigers. The Caspian tiger likely dispersed from an ancestral Northeast Asian population and experienced gene flow from southern Bengal tigers. Lastly, genome-wide monophyly supported the South China tiger as a distinct subspecies, albeit with mitochondrial paraphyly, hence resolving its longstanding taxonomic controversy. The distribution of mitochondrial haplogroups corroborated by biogeographical modeling suggested Southwest China was a Late Pleistocene refugium for a relic basal lineage. As suitable habitat returned, Eastern China became a genetic melting pot to foster divergent lineages to merge into South China tigers and other subsequent northern subspecies to develop. Genomic information retrieved from ancient tigers hence sheds light on the species’ full evolutionary history leading to nine modern subspecies and resolves the natural history of surviving tigers.

## Introduction

The radiation of modern Felidae began with the divergence of an ancient lineage that eventually gave rise to today’s five roaring cat species^1^, specifically the lion (*Panthera leo*), jaguar (*P. onca*), snow leopard (*P. uncia*), leopard (*P. pardus*), and tiger (*P. tigris*). Unlike some congeneric species that spread out to become the apex predators in other continents, the tiger originated, evolved, and remained in the jungle of Asia^1, 2^. Fossil records of probable *P. tigris* dated back to the early Pleistocene at c. 2.0 million years ago (Mya) from northern China and Java (Indonesia)^3–7^. Despite its once widespread presence, the glacial and climate oscillations during the Middle to Late Pleistocene likely caused the tiger range to repeatedly contract and expand. This resulted in a prolonged demographic bottleneck^5, 8–10^ that can be seen in their low genomic diversity and a mtDNA time to the most recent common ancestor (TMRCA) of only approximately 110,000 years ago^11–16^.

It is possible that waves of expansions after the Late Pleistocene bottleneck, followed by isolation across a heterogeneous landscape, have resulted in the present biogeographical pattern of tigers^9, 10^. Historically nine tiger subspecies inhabited a vast region west to the Caspian and Aral Seas, east to Northeast Asia, and south to the Sunda Islands^5, 8^ (see Table 1 for names and codes). During the last century, however, habitat loss, fragmentation, and hunting reduced the number of free-ranging tigers from probably over 100,000 to fewer than 4,000 today^17^. The Bali (BAL, *P. t. balica*), Caspian (VIR, *P. t. virgata*), and Javan (SON, *P. t. sondaica*) tigers are all extinct and the South China tiger (AMO, *P. t. amoyensis*) has not been seen in the wild for more than 30 years^17–19^. Recent genome-wide evolutionary analyses confirmed highly restricted gene flow across the range and supported six living subspecies, namely, the Amur (ALT*, P. t. altaica*), Indochinese (COR, *P. t. corbetti*), Malayan (JAX, *P. t. jacksoni*), Sumatran (SUM, *P. t. sumatrae*), Bengal (TIG, *P. t. tigris*), and South China tigers (AMO) ^16, 20^. Signals of selection have also been revealed in several subspecies, likely associated with local adaptation to various ecosystems^15, 20^.

**Table 1.** Ancient and modern tiger samples of the study including all nine modern subspecies and ancient Russian Far East population.

| Scientific name | Common name | Abbreviation | Sample quality | Sample age | Sample size | Number of mitogenome retrieved | Number of WGS retrieved | Source |
| --- | --- | --- | --- | --- | --- | --- | --- | --- |
| *Panthera tigris* | Ancient RFE tiger | RUSA | Zooarchaeological/<br>Ancient | 1,000~10,000 years | 25 | 7 | 1 | This study |
| *P. t. amoyensis* | South China tiger | AMO | Historical | ~100 years | 31 | 12 mitogenomes and 16 partial mtDNA sequences | 12 | This study |
|  |  |  | Fresh | Modern | 1 | 1 | 1 | This study |
|  |  |  | Fresh | Modern | 1 | 1 | 1 | <sup>20</sup> |
| *P. t. virgata* | Caspian tiger | VIR | Historical | ~100 years | 6 | 6 | 3 | This study |
| *P. t. sondaica* | Javan tiger | SON | Historical | ~100 years | 1 | 1 | 0 | This study |
| *P. t. balica* | Bali tiger | BAL | Historical | ~100 years | 1 | 1 | 0 | This study |
| *P. t. sumatrae* | Sumatran tiger | SUM | Fresh | Modern | 7 | 7 | 7 | <sup>20</sup> |
| *P. t. corbetti* | Indochinese tiger | COR | Fresh | Modern | 6 | 6 | 6 | <sup>20</sup> |
| *P. t. jacksoni* | Malayan tiger | JAX | Fresh | Modern | 9 | 9 | 9 | <sup>20</sup> |
| *P. t. altaica* | Amur tiger | ALT | Fresh | Modern | 6 | 6 | 6 | <sup>20</sup> |
| *P. t. tigris* | Bengal tiger | TIG | Fresh | Modern | 3 | 3 | 3 | <sup>20</sup> |

Several questions remain if the natural history of the tiger is to be fully resolved. Firstly, the contrast between the species’ age and its mtDNA coalescence indicates that the genomic diversity and demographic dynamics of ancient tigers might differ from that of their post-bottleneck counterparts. Secondly, while phylogenetic analysis of modern tigers clustered the Sumatran (SUM), Javan (SON), and Bali (BAL) tigers into a monophyletic clade, supporting a single ancient colonization event to the Sunda Island^12^, in contrast, the evolutionary pathways of mainland tigers are more complicated. For example, mtDNA gene sequencing suggested the extinct Caspian tiger is nearly indistinguishable from the Amur tiger^11^. A craniometric analysis showed an extensive overlap between the Caspian tiger and the other mainland subspecies^21^, implying an unsettled origin of the Caspian subspecies. Finally, previous population genomic analysis of modern specimens revealed a mtDNA lineage in South China tigers that is basal to all other subspecies and another divergent lineage similar to Indochinese tigers ^16, 20^. Whether this paraphyly is due to an admixed ancestry in the origin of South China tigers or a poorly defined geographic boundary between Indochinese and South China tigers would require an explicit assessment of voucher specimens of confirmed heritage^9^. Overall, it is likely that genomic scale data from ancient samples will be key for helping build a complete understanding of the evolution of tigers.

The past decade has witnessed substantial advances in ancient DNA research, now enabling the retrieval of genomic information from extinct mammals that date back to over one million years ago^22–24^, including Felidae such as saber-toothed cats^25–27^, the Late Pleistocene Holarctic lions^28–31^, and the European Late Pleistocene leopards^32^. However, thus far, only one whole genome^33^ has been retrieved from Pleistocene/early Holocene tiger specimens, partially due to the foregoing overlap of the tiger distribution with temperate-tropical forest biomes, where specimens are less likely to be preserved than in colder regions.

To elucidate the more comprehensive evolutionary history of tigers from the Late Pleistocene to present, we collected and generated high-quality genomes from over 60 samples spanning a wide geographic range and time scales, including both zooarchaeological/ancient (i.e., 1,000 ∼ 10,000 years) remains excavated from Northeast Asia and historical (i.e., ∼ 100 years) museum specimens that originated in Central and East Asia (Table 1, Fig.1, Tables S1, S2). These samples, including one from the Russian Far East (RFE) dated at c. 10,600 BP and the South China tiger holotype specimen designated in 1905^34^, are crucial for understanding tiger dispersal and inhabitation across mainland Asia. Together with 32 published genomes from extant subspecies, we provided an illuminating picture of the tiger’s evolutionary history, resolved the controversy in phylogeny and taxonomy, revealed cryptic refugia in mainland China, and contributed to the ultimate understanding of tiger genomic diversity that may guide the world’s conservation efforts of this iconic endangered species.

**Fig. 1.**
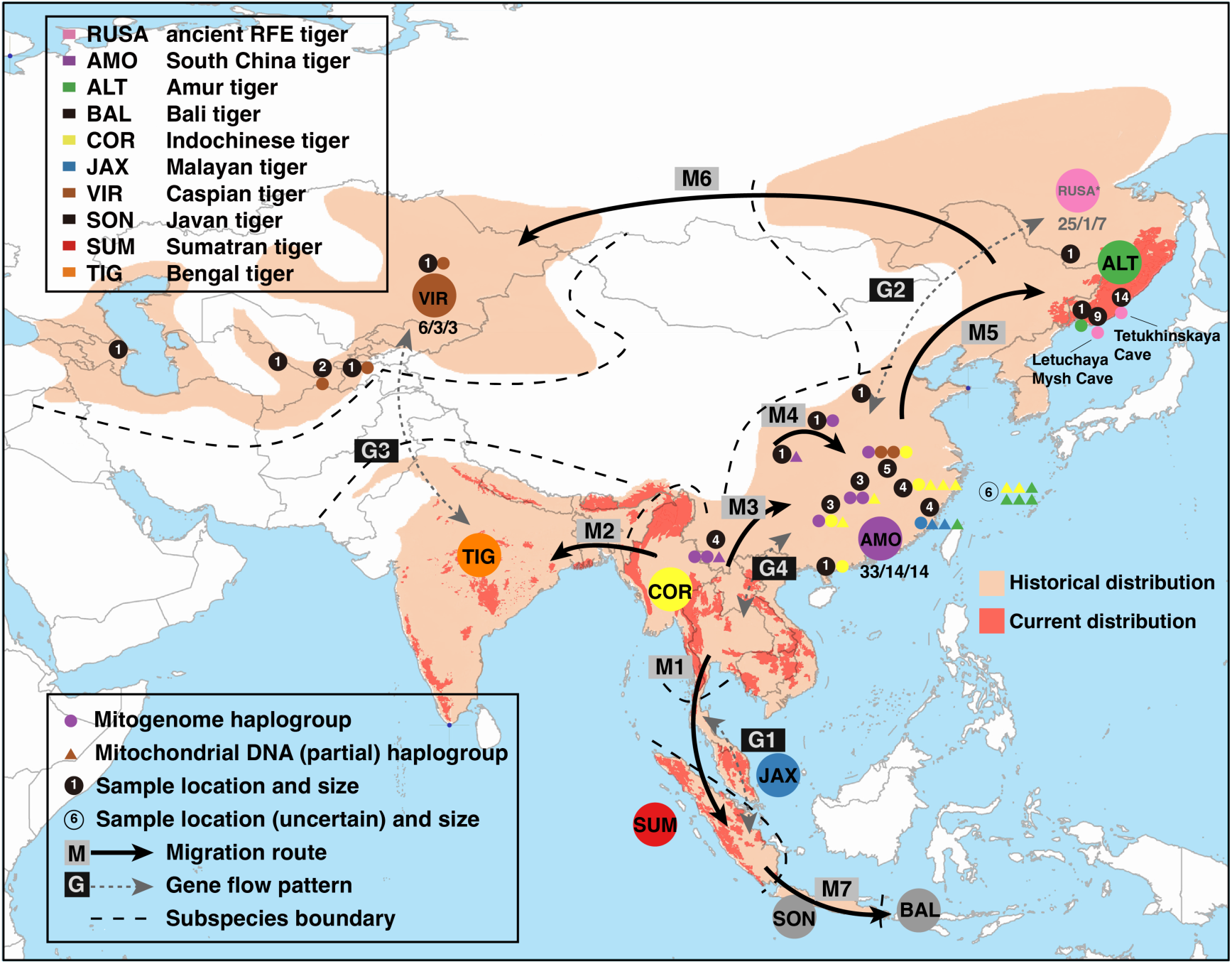
Ancient tiger specimens and genetic information collected in this study from the tiger’s northern range including China (N=33), the Russian Far East (RFE, N=25), and Central Asia (N=6) and the inferred dispersal routes of the species based on joint interpretation from modern and ancient tiger genomes. Small circles with numbers indicate the geographic origin and sample size from each location. Large colored circles with abbreviations correspond to the distribution of the nine modern subspecies and the ancient RFE population. The evolutionary scenarios were reconstructed based on the genome-wide phylogeny, Bayesian molecular dating, modeling of migration and gene flow, and ecological niche modeling of suitable habitats. Labels (x/y/z) on the subspecies codes of the South China tiger (*P. t. amoyensis*, AMO), the Caspian tiger (*P. t. virgata*, VIR), and the ancient RFE tigers (RUSA) correspond to total sample size/the number of whole-genome datasets acquired/the number of mitogenomes assembled. The maternal ancestry of each sample from China and the RFE is color coded according to its association with modern subspecies or ancient populations and based on either mitochondrial genome (small, solid, colored circles) or partial mitochondrial DNA (small, solid, colored triangles) sequencing.

## Results

### Genome dataset of extinct tigers

We retrieved genome sequences from seven of the 25 ancient RFE felid fossil remains and evaluated their endogenous DNA authenticity (Table S1, Fig. S2). The final ancient RFE tiger genome dataset included one whole-genome assembly with 8× coverage from a phalanx bone (RUSA21) excavated from the Letuchaya Mysh Cave and six mitogenomes obtained by RNA probe target capture enrichment (Table 1, Fig. 1). RUSA21 was radiocarbon dated to c. 10,600 BP and other six specimens were dated to c. 8,600-1,800 BP (Table S1).

We obtained genome sequencing data over timescales of centuries from tigers recently extirpated from the range, including South China tigers (AMO, *P. t. amoyensis*) and Caspian tigers (VIR, *P. t. virgata*), Javan tigers (SON, *P. t. sondaica*), and Bali tigers (Bali, *P. t. balica*) (Fig.1, Tables 1 and S2). Eleven South China tiger specimens were sequenced to 1-12×, including one holotype (museum ID 3311) and three paratypes (3305, 3307, 3308) collected in Hankou, Hubei Province, China^34^.

Three Caspian tiger specimens originated from Uzbekistan and Kazakhstan were sequenced to 4-8×. We also sequenced 16 South China tiger museum specimens with partial mtDNA fragments encompassing subspecies-diagnostic sites (Fig. 1, Table S2), assembled two mitogenomes using next-generation sequencing data from Javan and Bali tigers, and incorporated these data into the phylogenetic analysis. The geographic distribution of these samples hence fully covered the historical range of the extinct tiger subspecies. In combination with 32 previously published whole-genome sequences^20^, our final dataset consists of 48 nuclear genomes including 15 newly produced genomes from ancient specimens and 39 mitogenomes including 17 novel ones.

### Genome-wide phylogeny and taxonomy of living and extinct tigers

A whole-genome phylogeny was reconstructed based on over 1.2 million transversions that were identified from the autosomal neutral regions of 15 ancient and 33 modern tiger genomes. In addition to those of the five subspecies that were highly distinct as reported before^16, 20^, the newly assembled genomes of South China tiger and Caspian tiger voucher samples each formed a monophyletic group (Fig. 2), thus providing robust genome-wide evidence consistent with the subspecies classification and taxonomic designation of modern tigers. The Sumatran Island subspecies *P. t. sumatrae* (SUM) was the earliest branch in the species’ phylogeny. Within mainland Asia, one monophyletic clade consisting of all three Bengal tigers (TIG) is sister to another including all Caspian tigers (VIR), although bootstrap support for the TIG-VIR grouping was not statistically significant (Fig. 2A). The 10,600-year-old RFE tiger RUSA21 clustered closely to present-day Amur tigers (ALT) from the same region, and together they formed the sister clade to South China tigers, a monophyletic AMO group including nine of the 11 South China tigers. The last major clade of tigers comprised the phylogenetically robust Malayan tigers (JAX) and a paraphyletic assembly of Indochinese tigers (COR). Two South China tiger (AMO) specimens (Y14 and NGH2) originated from Southwest China clustered as an outgroup to the Southeast Asian monophyletic clades (COR and JAX, Fig. 2A). For WGS analysis, Y14 and NGH2 are the only exception to the monophyly of the tiger subspecies, likely reflecting the gene flow between adjacent Indochinese and South China tigers in their contact zone.

**Fig. 2.**
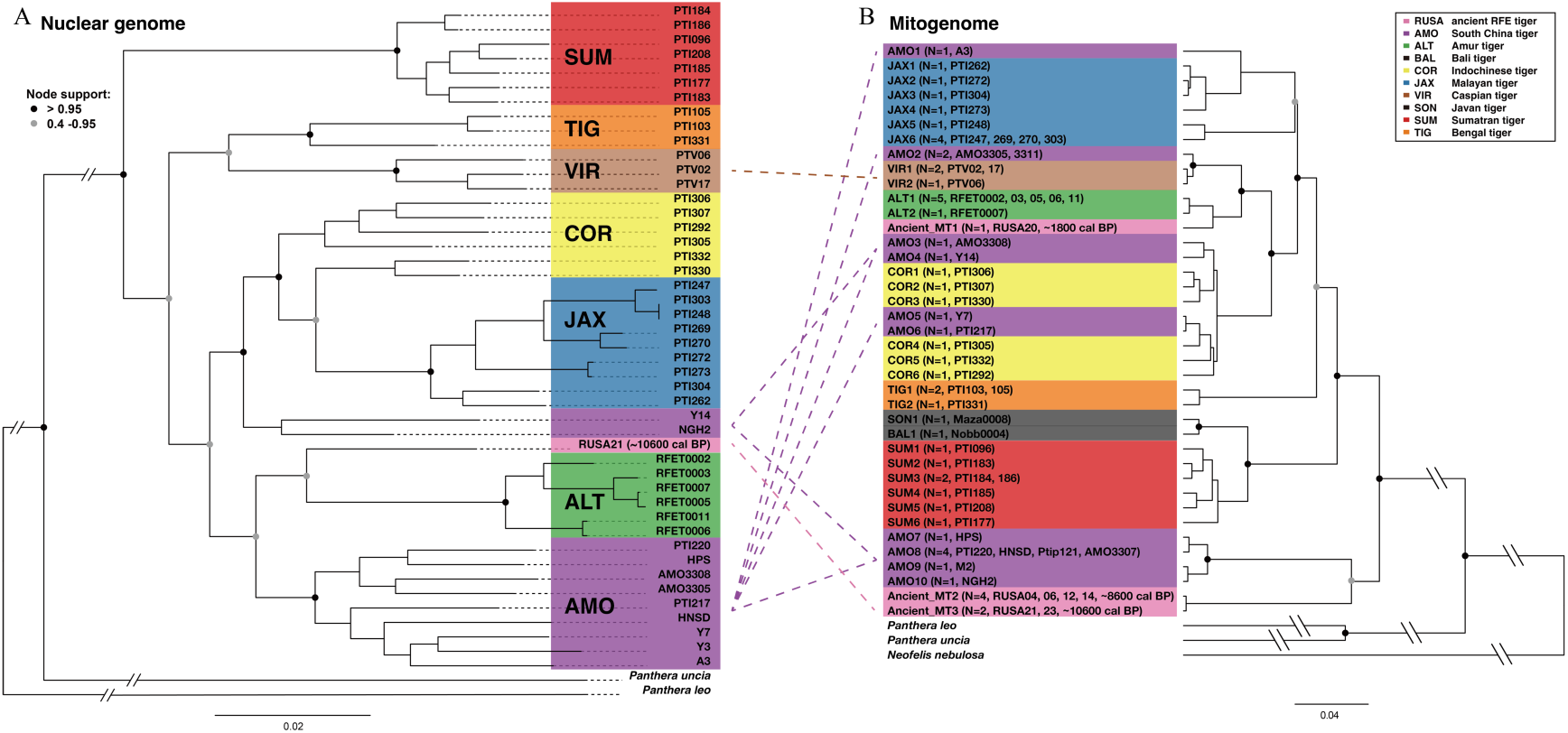
Phylogenetic relationships among ancient and modern tigers. Sample labels are color-coded according to their taxonomic classifications based on geographical origin. Clades with high bootstrap support are marked on major nodes. (*A*) Bayesian phylogeny inferred from 15,450 bp of mitogenome sequences excluding the control region. Sample labels are mtDNA haplotype codes, followed by the number of samples sharing the haplotype and sample IDs in parentheses. Mitogenome phylogenetic trees based on neighbor-joining (NJ), maximum parsimony (MP), maximum likelihood (ML), and Bayesian analyses give identical topologies, and the Bayesian tree is shown here. Nodes with posterior probabilities of 0.4-0.95 and over 0.95 are indicated. (*B*) ML phylogeny inferred from 1,242,142 transversions in genome-wide autosomal neutral regions. Node support was evaluated with 100 bootstrap replicates. Samples are labeled at the tree tips, and the same ancient or historical specimens in the two phylogenetic trees are indicated by dotted lines showing cytonuclear discordance in the South China tiger (*P. t. amoyensis*), the Caspian tiger (*P. t. virgata*), and the ancient RFE tiger (RUSA21).

A phylogenetic tree based on 15,450 bp of mitogenome sequences (excluding the control region) exhibited cytonuclear discrepancies in South China tiger (AMO), Caspian tigers (VIR), and ancient RFE (RUSA) sampled from different geological time periods (Fig. 2B). In contrast to the Caspian tiger (VIR) *–* Bengal tiger (TIG) association indicated by the autosomal phylogeny, the mtDNA genealogy indicated no close relationship between the two subspecies. Two VIR mitogenomes and one mitogenome sequence AMO2 shared by two AMO paratype specimens (3305 and 3311) clustered together, which then formed the sister clade of Amur tigers (ALT, Fig. 2B).

We detected five mitogenome haplogroups among the 14 South China tiger (AMO) voucher specimens that respectively aligned with those found in the Malayan tiger (JAX, N=1), Caspian tiger (VIR, N=2), two Indochinese tiger lineages (COR, N=4, including AMO paratype 3308), or formed a distinct clade basal to all extant tiger subspecies (N=7). The AMO holotype specimen (3311), collected from Hankou of Hubei Province in East China, was most closely associated with Caspian tiger. In addition, we retrieved partial mtDNA sequences from 16 historical specimens of South China tigers and identified their maternal ancestry based on subspecies-diagnostic sites^20^ (Table S2, Fig. 1). The mtDNA lineages of these samples belonged to the Malayan tiger (JAX, N=2), Amur/Caspian tigers (ALT/VIR indistinguishable from the partially sequenced mtDNA fragment, N=5), Indochinese tiger (COR, N=7), and the basal AMO lineage uniquely found in South China tigers (N=2).

In sharp contrast to the autosomal monophyly, the 30 South China tigers mtDNA haplotypes consistently showed paraphyly - even within the four type specimens. Since all samples were traceable to geographic origins within the generally recognized South China tiger range, we hypothesized that the cytonuclear discordance is most likely an intrinsic feature of the natural population, rather than artificially introduced admixture in captivity as previously suggested^16, 20, 35^. Interestingly, the AMO mtDNA haplotypes shared with other subspecies were restricted to the eastern part of China but absent in the western regions, such as Shaanxi, Yunnan, and Guizhou, where only the unique basal AMO mtDNA lineages were present (Fig. 1). Despite mitochondrial DNA paraphyly in South China tiger, autosomal DNA monophyly rendered strong support for *P. t. amoyensis* as a statistically robust subspecies.

In the RFE region, the consistency of sequence identity, excavation location, and radiocarbon dating data across different samples supported that the seven ancient RFE tiger specimens (RUSA) might belong to three individuals or maternal lineages (Table S1, Table S2, and Fig. 1). The mitogenome sequence of RUSA23 was identical to that of RUSA21 (Ancient_MT3) from the Letuchaya Mysh Cave, dated to c. 10,600 BP. Four tooth or bone specimens from the Tetukhinskaya Cave (RUSA 04, 06, 12, 14) shared the same mtDNA haplotype (Ancient_MT2), all of which were dated to c. 8,600 BP. Ancient_MT2 and Ancient_MT3 consisted of the sister clade of the unique South China tiger haplogroup, together forming the basal linage of all modern tigers. RUSA21 was most closely related to the living RFE tiger populations according to the autosomal data, a pattern not evident in the mitochondrial phylogeny. In contrast, RUSA20, a more recent specimen from the Letuchaya Mysh Cave, was dated to c. 1,800 BP and carried a mitogenome haplotype (Ancient_MT1) closely associated with that of extant Amur tigers. The three ancient lineages spanned a chronology of 10,000 years and recapitulated the evolutionary turnover of tigers on the northeastern edge of the range.

### Population genomic structure

Analyses of population genetic structure based on the same dataset of 48 extinct and extant tiger genomes provided further support for the partitioning of *P. tigris* subspecies (Fig. 3). The first two principal components (PC1 and PC2) in the principal component analysis (PCA) explained approximately 20% of the total variance and distinguished Sumatran (SUM), Malayan (JAX), and Amur (ALT) tigers. PC3 and PC4 accounted for an extra 10% of the total variance and segregated Bengal (TIG) and Indochinese (COR) tigers. The newly sequenced South China (AMO) and Caspian (VIR) tiger specimens clustered within themselves, although not with the distinctiveness as seen in other subspecies (Fig. 3A), likely reflecting the existence of historical gene flow during the evolutionary past. Consistent with the cytonuclear phylogenetic discordance, the ancient RFE tiger RUSA21 was closely associated with South China tigers in the PCA, not with modern Amur tigers from the same region.

**Fig. 3.**
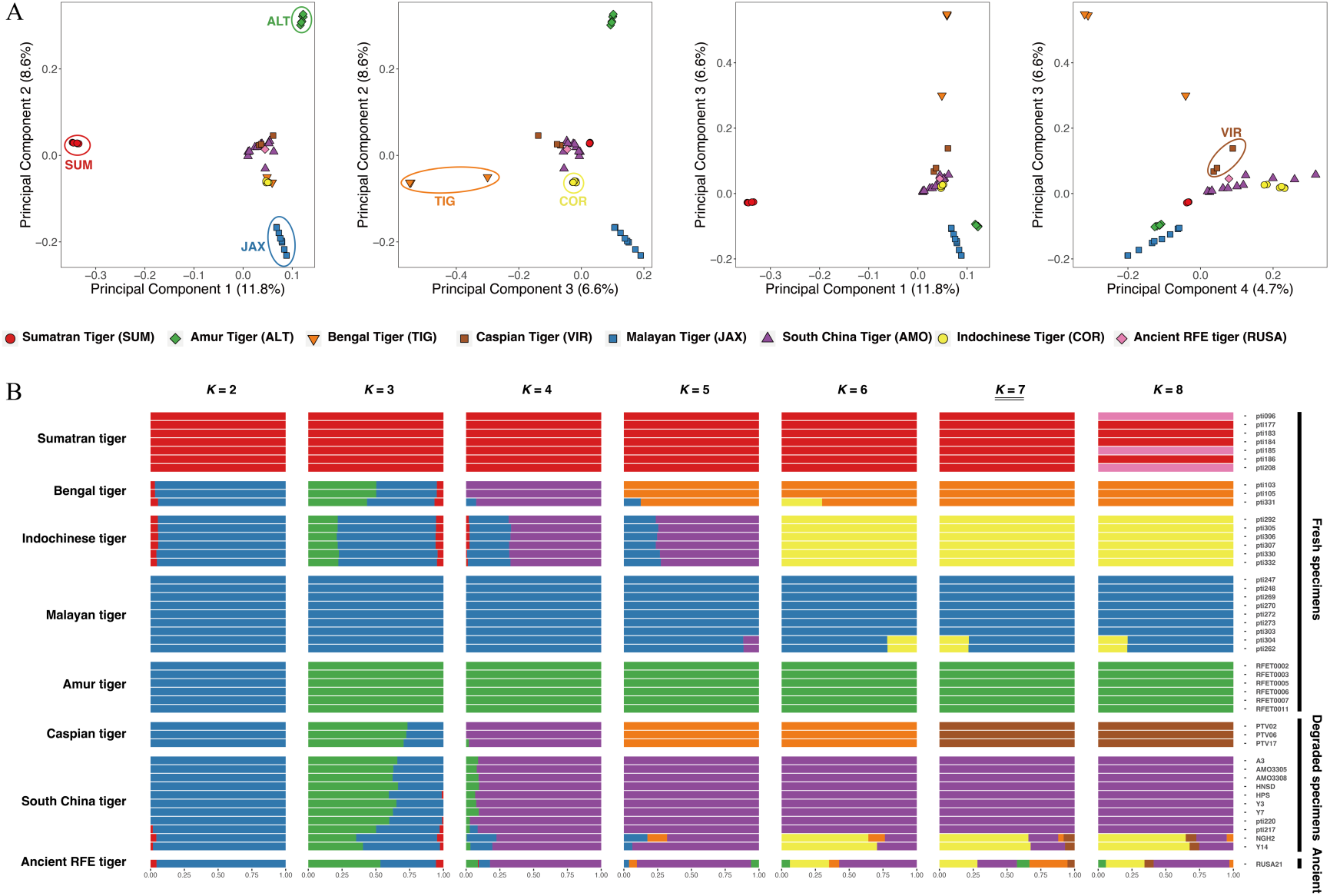
Population genomic structure of 13 ancient and 33 modern tigers based on 1,242,142 transversions from autosomal neutral regions. (*A*) Principal component analysis (PCA) of ancient and modern tigers showing the first four principal components. Each tiger is represented by one icon whose color and shape correspond to the subspecific affiliation of the specimen. (*B*) Population genetic structure estimated in ADMIXTURE with the number of inferred populations (*K*) from 2 to 8. Each horizontal bar corresponds to one tiger individual and is proportionally partitioned into *K* colored segments representing the individual affiliations with each cluster. When *K*=7, the seven inferred ancestral populations are in accordance with the seven highly distinctive modern tiger subspecies. Two *P. t. amoyensis* and the ancient RFE tiger show admixed genomic compositions.

Population structure assessment in ADMIXTURE also clustered the tiger genomes according to their subspecies affiliation when the number of groups (K) was set to 7 (cross-validation error (CVE) = 0.79, Fig. 3B). All Caspian tigers and all but two South China tigers displayed their own population distinctiveness. The same two tigers from Southwest China (Y14 and NGH2) that fell within the Indochinese tiger clade in the autosomal phylogeny also showed over 60% genetic identity with Indochinese tigers. RUSA21 possessed a highly admixed background including genetic components from nearly all tiger subspecies across mainland Asia, albeit with the largest proportion from South China tigers. Similar patterns shown by the PCA, ADMIXTURE, and phylogeny results suggested a likely connection between the modern South China tiger (AMO) and the ancient RFE tiger population 10,000 years ago.

### Ancient gene flow scenarios

To disentangle whether gene flow may have contributed to the cytonuclear discordance revealed by the phylogenies, *D*-statistics were applied to evaluate for possible ancient admixture scenarios based on the excessive amount of allele sharing between populations. Post-divergence gene flow was evident among some subspecies of geographical proximity, for instance, between the Sumatran tiger and the ancestral population from which Indochinese and Malayan tigers were derived (Fig. 4A). Genetic admixture of Indochinese and Malayan tigers with other populations likely also reflected the ancient events prior to the splitting of the two subspecies (Fig. 4B).

**Fig. 4.**
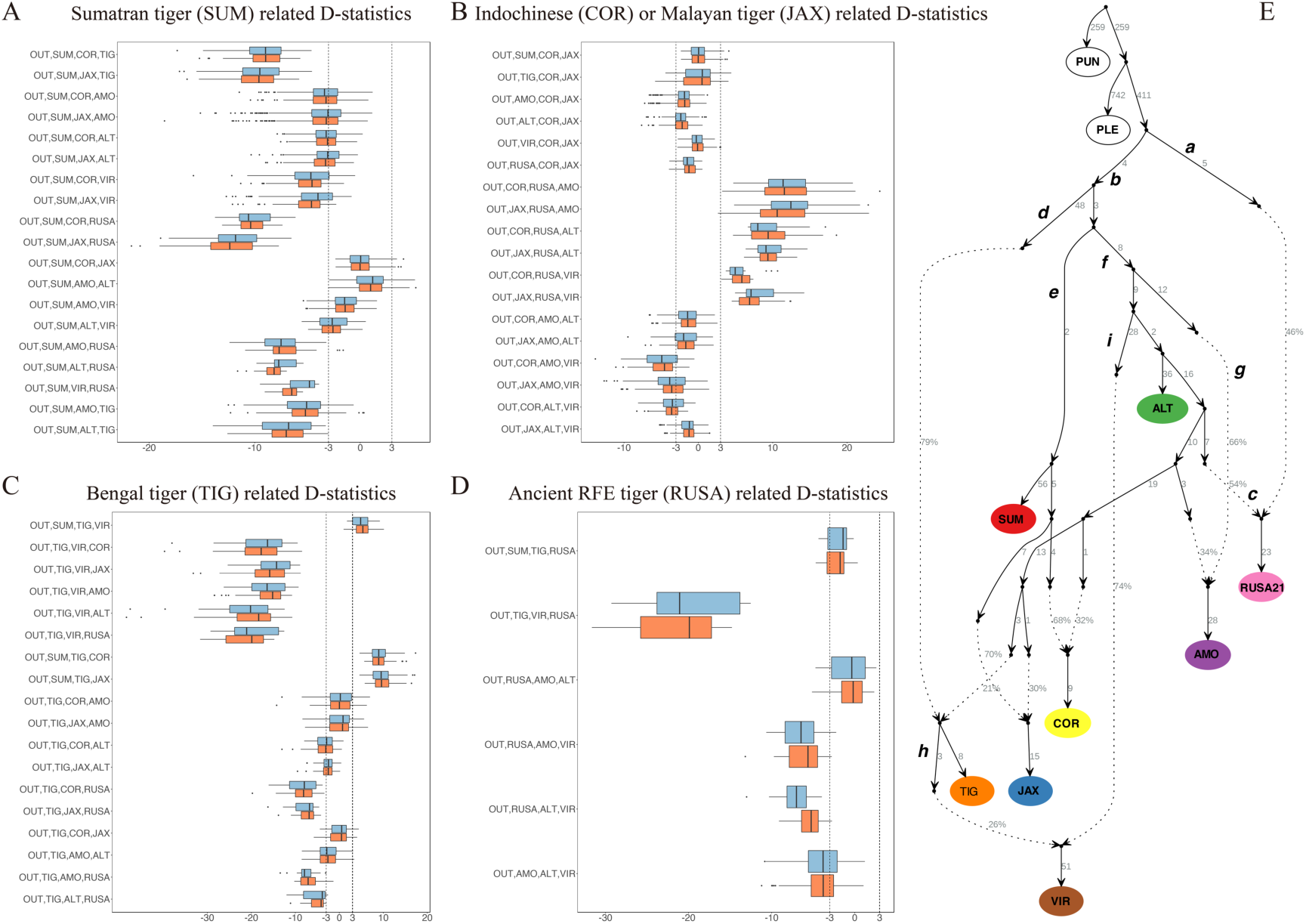
*D*-statistic inference of the genetic ancestry of tigers and *f*-statistic-based admixture graph modeling based on ancient and modern specimens. (*A-D*) *D*- statistics testing for possible scenarios of gene flow related to different tiger populations. Transformed z-scores are shown, with |z| > 3 considered significant, in which z > 3 means that the extent of allele sharing between “Pop” and “Pop2” is greater than that between “Pop” and “Pop1” for the *D* (Outgroup, Pop; Pop1, Pop2) setting, while z < -3 might reflect genetic admixture between “Pop” and “Pop1”. The color of the boxes refers to the different tests in which the lion (orange) or snow leopard (blue) is used as the outgroup. (*E*) Admixture graph modeling of the ancestry of tigers with the best-fitting model shown. Solid lines represent phylogenetic relationships among tiger populations, and numbers mark branch length in units of genetic drift. Dotted lines show the inferred scenario of genetic admixture events between lineages, with the percentages corresponding to admixture proportions from two ancestral populations. Admixture graphs were built using transversion sites only, and one sample with the highest sequencing depth was used to represent each subspecies. Abbreviations for the specimens are as follows based on the geographic origin of the individual: AMO, the South China tiger (*P. t. amoyensis*); ALT, the Amur tiger (*P. t. altaica*); COR, the Indochinese tiger (*P. t. corbetti*); JAX, the Malayan tiger (*P. t. jacksoni*); SUM, the Sumatran tiger (*P. t. sumatrae*); TIG, the Bengal tiger (*P. t. tigris*); VIR, the Caspian tiger (*P. t. virgata*); RUSA, the ancient Russian Far East tiger population dated to approximately 10,000 years ago; PLE, the lion *P. leo*; PUN, the snow leopard *P. uncia*.

The amount of allele sharing between a modern subspecies and the ancient RUSA21 was consistently minor in comparison to that between any two modern subspecies (Fig 4), indicative of a evolutionary turnover of RFE tiger populations over the past 10,000 years. It is worth noting however, that the amount of allele sharing between RUSA21 and the South China tiger (AMO) or Amur tiger (ALT) was higher than that between RUSA21 and the Caspian tiger (VIR, Fig. 4D). This pattern may reflect a close relationship between the 10,600-year-old RFE tiger population and the ancestral population in East Asia from which modern South China and Amur tigers were later derived. It also supported an absence of post-divergence gene flow between this ancient East Asian population and the westbound Caspian tiger. In contrast, historical gene flow was significant between Bengal (TIG) and Caspian (VIR) tigers, which, together with phylogenetic and population clustering patterns, indicated an evolutionary mechanism underlying the cyto-nuclear phylogenetic discrepancy (Fig. 4C).

The ancient admixture scenarios revealed by the D-statistics were integrated in *f*- statistics-based admixture graphs to reconstruct the genetic relationships between different extant and extinct tiger groups (Fig. 4E). The admixture graph allowing for gene flow events illustrated two major branches after tigers split from the *Panthera* outgroup, including one extinct “ghost” lineage (a in Fig. 4E) that contributed to approximately half of the genomic ancestry of the ancient RFE tiger (RUSA) and one phylogroup (b) comprising all extant tiger subspecies. The other half of RUSA21’s genome was from genetic mixture with another ancestral lineage in Northeast Asia (c) that was associated with today’s Amur (ALT) and South China (AMO) tigers.

The subsequent divergence leading to modern tiger subspecies can be further divided into three waves, with short relative branch lengths in *qpGraph* that are indicative of a rapid and nearly simultaneous radiation. One lineage (d) eventually formed the Bengal tiger (TIG), one (e) became the ancestor to the Sumatran tiger (SUM) and likely other tigers on the Sundaland, and the third (f) established the other subspecies on mainland Asia including an early split (g) leading to the South China tiger (AMO). Both admixture graph modeling (Fig. 4E) and TreeMix phylogenetic approaches, which considered different migration bands (from 1 to 7, Fig. S2), supported a model where the admixed ancestry of the Caspian tiger (VIR) in Central Asia derived from (i) an ancient lineage (h in Fig. 4E) related to the Bengal tiger and (ii) a Northeast Asian lineage (i in Fig 4E) ancestral to modern Amur (ALT) and South China (AMO) tigers. Overall, the demographic history concerning post-divergence gene flow demonstrated that most admixture events occurred before the latest divergences giving rise to modern subspecies.

### Coalescence dating and demographic modeling

The 10,600-year-old ancient RFE tiger specimen (RUSA) and the extinct Caspian (VIR) and South China (AMO) tigers showed a pattern similar to that of other extant subspecies in the pairwise sequential Markovian coalescent (PSMC) model (Fig. S3). All tigers experienced a decline in effective population size (*Ne*) during the Late Pleistocene that was likely related to the divergence of modern tiger subspecies caused by climate change.

The TMRCA of all tigers’ mitogenomes was traced back to 107,200 years ago (95% confidence interval (CI): 67,900-148,400 years ago) using a Bayesian method implemented in BEAST2 (Fig. 5A). Ancient RFE specimens from this study clustered within the modern tiger clade and hence did not alter the species’ matrilineal TMRCA estimates based exclusively on modern samples^12–14, 16, 20^. The two specimens aged to 8,600 BP and 10,600 BP were closely related and then distantly coalesced with the unique South China tiger lineage 91,900 years ago (95% CI: 53,900-130,700 years ago), corroborating their basal position in the mitochondrial phylogeny. The more recent specimen from 1,800 BP formed a clade with current Amur tiger samples from the same region, with the two mtDNA lineages shared a TMRCA at about 16,100 years ago (95% CI: 6,700-26,200 years ago). This pattern likely reflects the population continuity in the RFE over the past two millennia.

**Fig. 5.**
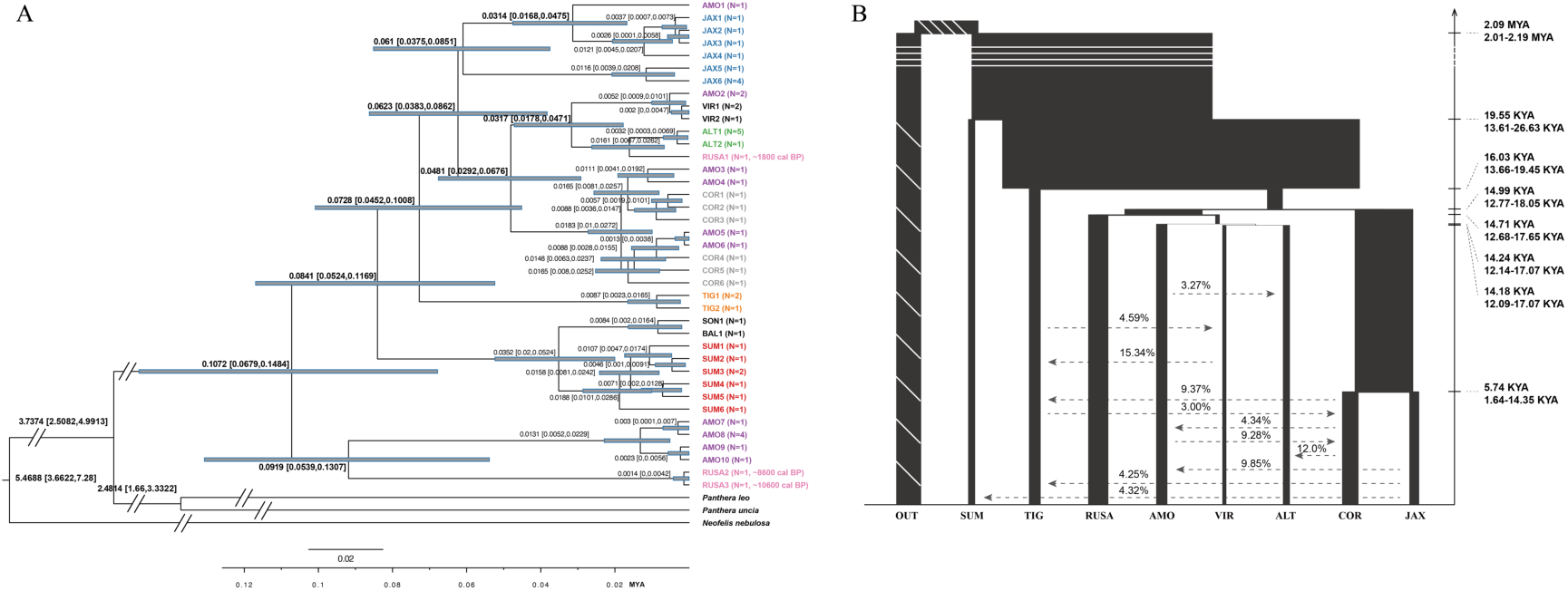
Population history modeling of tigers based on modern and ancient specimens. (*A*) Mitogenome divergence time estimated with BEAST2. Mitochondrial haplogroups are color-coded by the subspecies (modern tigers) or population (ancient Russian Far East tiger) source of the specimen, and the number of samples sharing the mtDNA haplotype are indicated in parentheses. (*B*) Demographic model inferred from G-PhoCS integrating the divergence time (proportional to the height of each bar), population size (proportional to the width of each bar), and total migration rate, with the direction indicated by arrowed lines. The three-letter code indicates the source of the specimen on the basis of geographic origin: AMO, the South China tiger (*P. t. amoyensis*); ALT, the Amur tiger (*P. t. altaica*); COR, the Indochinese tiger (*P. t. corbetti*); JAX, the Malayan tiger (*P. t. jacksoni*); SUM, the Sumatran tiger (*P. t. sumatrae*); TIG, the Bengal tiger (*P. t. tigris*); VIR, the Caspian tiger (*P. t. virgata*); RUSA, the ancient Russian Far East tiger population dated to approximately 10,000 years ago; OUT: *P. leo* or *P. uncia* as the outgroup.

G-PhoCS demographic modeling based on whole genome data (Fig. 5B) revealed the coalescent time of all tigers to be about 19,550 years ago (95% CI: 13,610-26,630 years ago). This estimate is substantially more recent than the estimated mitogenome coalescence, which is possibly due to subsequent gene flow after the initial population split. The oldest nuclear genome coalescent event occurred between the Sumatran tiger and mainland populations, followed by the earliest divergence within the mainland forming the Bengal tiger approximately 16,030 years ago (95% CI: 13,646- 19,450 years ago). The rapid radiation was most prominent in the northern part of the range, giving rise to the modern Amur tiger (ALT), Caspian tiger (VIR), South China tiger (AMO), and the ancient RFE tiger (∼10,000 years ago, RUSA). The final clear split of the Malayan tiger (JAX) from the Indochinese tiger (COR) was the latest, approximately 5,740 years ago (95% CI: 1,640-14,350 years ago). The total post-divergence migration between subspecies was low at 3-12% and mostly between geographically adjacent populations. The highest level of gene flow was inferred between the Bengal tiger (TIG) and Caspian tiger (VIR) at approximately 15%, in support of the contribution of the Bengal tiger to the genomic ancestry of the Caspian tiger. Due to the limitation of the parameter estimation in the complex model, all inferred migration bands were considered bidirectional.

### A historical tiger refugium in Southwest China

Phylogenomic patterns of mitochondrial paraphyly and autosomal monophyly in South China tigers (AMO), as well as demographic history and gene flow inference, jointly indicate a plausible scenario for resolving the longstanding controversy over the origin of *P. t. amoyensis*. The restricted distribution of the non-unique (i.e., shared with other subspecies) mtDNA haplotypes in eastern China, but not in the west, where only the basal lineage unique to the modern South China tiger was present (Fig. 1), prompted a hypothesis that Southwest China may have once been a refugium harboring relic tiger populations, while eastern China only became a genetic melting pot for various ancient lineages expanding into this region only as biogeographical conditions became optimal.

To test this hypothesis, we performed ecological niche modeling (ENM) to project tiger habitat suitability under the influence of climate fluctuation from the Last Interglacial (LIG, between c. 130,000 and 115,000 years ago) to the present (Fig. 6). The climate dynamics associated with the potential tiger range roughly corresponded to the alternating episodes of glacier advance and retreat. During mild and humid interglacial periods such as the LIG, Mid-Holocene, or present day (Fig. 6A, Fig. S6), suitable tiger habitat was widespread and continuous throughout much of Asia, from the Caspian Sea and the Indian subcontinent in the west, to Siberia in the north, to the Sunda Islands in the south, and to the Japanese archipelago in the Far East^10, 14^. During cool and dry periods such as the Last Glacial Maximum (LGM, approximately 22,000 years ago, Fig. 6B), although reduced sea level may have created land bridges to facilitate dispersal, substantial climate cooling likely exerted a profoundly negative impact on tiger landscapes, resulting in local extinctions and range fragmentations.

**Fig. 6.**
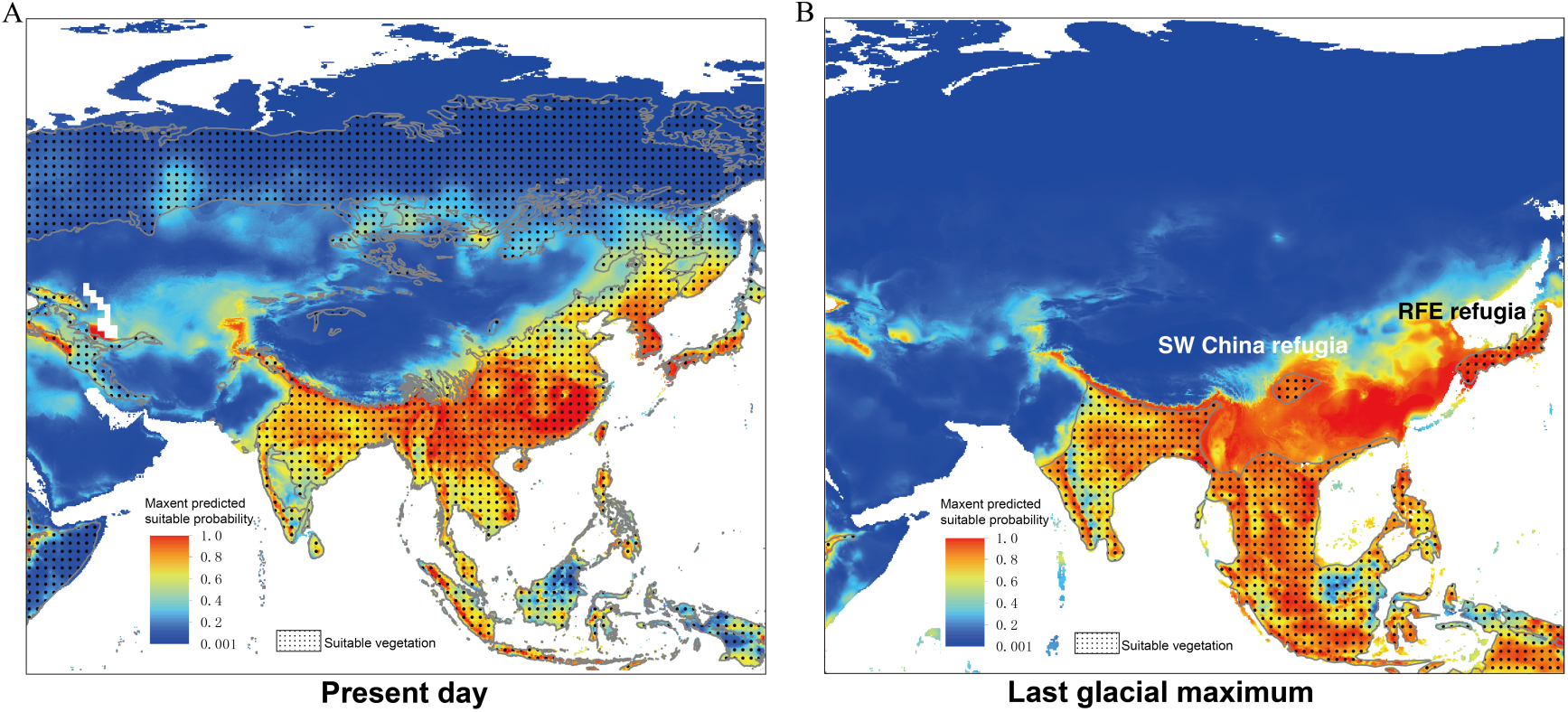
The projected tiger habitat suitability during the Last Glacial Maximum (LGM) and at present based on ecological niche modeling (ENM) in Maxent overlaid with the preferred vegetation layer. The Maxent model was built with present-day tiger occurrence data collected from the literature based on nine bioclimate variables. The probability of suitability for the LGM was projected based on climate variables during the LGM. Possible refugia with suitable habitat for the tiger during the LGM are highlighted in Southwest China (SW China) and the Russian Far East (RFE). Different colors refer to different levels of likelihood of tiger presence. The dotted regions represent vegetation zones suitable for tigers with data assembled from the IUCN Red List of Threatened Species for the present-day layer^83^ and from published studies for the LGM map^84^.

When ecological niche models were overlaid with the vegetation types suitable for tigers (Fig. 6), a further-restricted tiger distribution during an LGM-like period was apparent, in which the majority of today’s eastern part of China was composed of temperate grassland interspersed with sparse woodland and was not optimal habitat for tigers. In contrast, two possible northern Asian refugia that may have sustained tigers and prey populations were revealed, including one pocket zone in the mountains of Southwest China and another on the Japanese archipelago. Although the arrival and extinction of tigers in ancient Japan remained somewhat elusive^36–38^, the scenario of a refugium in Southwest China concurrent with the suboptimal habitat in the east during an LGM-like period supported the cyto-nuclear discrepancy observed in South China tigers and a post-glacial genetic melting pot hypothesis in China.

### Genetic melting pot in eastern China during tiger expansion

Multiple genetic studies of the tiger^15, 16, 20^ converged on a low level of genomic diversity that corresponded to a Late Pleistocene demographic bottleneck in the species. TMRCA for mitogenome suggested that this prolonged contraction was likely associated with the beginning of the Last Glacial Period approximately 115,000 years ago. Subsequently, this prehistoric population underwent a modest growth and an overall decline, when it differentiated into multiple phylogroups associated with various habitats and selection pressures (Fig. S3).

At the advent of a glacial period, driven by the dry and cold climate, the connection between some tiger populations across mainland Asia might have disappeared and relic lineages would have become scattered across the fragmented landscape. One ancestral lineage carrying the unique South China tiger mitochondrial haplotype probably survived in the mountainous Southwest China. The long-lasting isolation ended after the glacial maximum. The post-bottleneck radiation center was likely located in the Indochina region and started with mainland–Sundaland isolation (Fig. 1, M1) followed by the split of the Bengal tiger to the Indian subcontinent (Fig. 1, M2). With a warmer climate and return of habitat across the Northern Hemisphere especially at the high latitudes, the ancient populations in Indochina rebounded and dispersed into eastern China (Fig. 1, M3), where they encountered the relic tiger population expanding out of the Southwest China refugium (Fig. 1, M4). The eastern part of China, then covered with habitats suitable for tigers such as forest and shrubland, may have functioned as a “melting pot” where distant and previously isolated lineages merged to create the genetic makeup of contemporary South China tigers (*P. t. amoyensis*).

After the last glacier completely retreated within the past 12,000 years, tigers in eastern China continued their dispersal to the north and reached Northeast Asia, probably in multiple waves driven by climate fluctuations. One of the earlier waves likely involved an extinct lineage represented by the 10,600-year-old tiger specimen RUSA21 and the latest turnover gave rise to the modern Amur tiger (M5 in Fig. 1). The Caspian tiger in Central Asia originated from two sources, first established from the expansion of an ancestral Northeast Asian population through the Siberian corridor (Fig. 1, M6) and then admixed with a northern-bounded population from the Indian subcontinent (Fig. 1, G3). The divergence of modern tiger subspecies began with the isolation driven by climate fluctuations during the last glacial period and was completed after the last wave of migration after the LGM.

## Discussion

In this study, genomic analysis of tiger specimens spanning the past 10,000 years revealed the most comprehensive evolutionary history for the species to date. Distinctive population structure and genome-wide phylogenetic monophyly supported the South China tiger (*P. t. amoyensis*) and Caspian tiger (*P. t. virgata*) as statistically robust subspecies, yet cyto-nuclear discrepancy also revealed genetic admixture from other lineages contributing to the origin of the subspecies. Genomic ancestry, demographic modeling, and historical habitat reconstruction suggested multiple tiger range contraction–expansion–isolation cycles in northern mainland Asia that were likely associated with the glacial cycle and climate fluctuations. Such patterns led to the proposed scenarios of the cryptic tiger refugia in Southwest China^39, 40^, genetic homogenization of various distinct lineages in eastern China following the return of habitat, and multiple waves of population expansion, admixture, and turnover in the RFE area.

The 10,600-year-old tiger specimen (RUSA21) excavated from the RFE, from which we produced a high-quality ancient genome with 8.1× coverage, possibly represented one of the earlier post-LGM waves of tiger expansion to the northeastern edge of the species’ range, based on ENM (Fig. S5). Fossil records of probable tigers date back to the early Pleistocene at c. 2.0 Mya^4, 6, 7^, and possibly to the Pliocene – Pleistocene boundary at c. 2.6–2.2 Mya if a tiger-like felid, *Panthera zdanskyi*^6^, is considered a tiger. However, the mtDNA coalescence time of modern tigers was traced back to a maximum of 110,000 years ago^15, 16, 20^. Such a contrast between the species’ age and TMRCA suggests a prolonged demographic bottleneck in ancient tigers, whose evolutionary genomic pattern might differ from that of modern tigers and harbor genetic diversity lost from contemporary samples. Our results, which focus on the recent evolutionary history of the tiger since the TMRCA, support this hypothesis in terms of the genetic ancestry of RUSA21, which involved an extinct and early-diverging lineage basal to and no longer present in modern tigers (Fig. 4E) as well as contribution from lineages related to extant Amur tigers (Fig. 2). ENM-based habitat modeling also projected the presence of refugia in Northeast Asia during a glacial maximum period that may have sustained relic tiger lineages. The likely presence of a refugium in the south of the RFE is evidenced by the findings in Late Pleistocene cave deposits of new endemic species of ochotonids and giant flying squirrels^41, 42^, as well as species that went extinct in the rest of the territory as early as the Early Pleistocene^43, 44^.

Our sampled RFE tiger specimens spanned different geological periods at c. 10,600 BP, c. 8,600 BP, c. 1,800 BP, and the present, and highlighted major temporal population turnovers in Northeast Asia. We discovered four mitochondrial haplogroups that corresponded to the four sampling periods and belonged to two clusters. Two mitogenomes, of which one was carried by two specimens from c. 10,600 BP and the other was shared by four from c. 8,600 BP, grouped with the unique South China tiger lineages to form the basal clades in the tiger mitochondrial phylogeny. In contrast, the recent specimen from c. 1,800 BP aligned with modern Amur tigers instead. The population turnover between c. 8,600 BP and c. 1,800 BP suggested possible replacement of the older lineages with more recent lineages showing migratory waves from eastern and northern China. Morphological studies of the Late Pleistocene tiger fossils also supported multiple waves of dispersal from the south to the Far East^33, 45^. On the other hand, the evolutionary continuity of RFE tigers from c. 1,800 BP to the present provided evidence for the probable establishment of the current Amur tiger subspecies by approximately 2,000 years ago.

We also elucidated for the first time the evolutionary genomic history of the tiger in its westmost range, featuring the establishment of the Caspian tiger (VIR). The extinct Caspian tiger once occupied a vast area from the riverine and forest habitat in Northwest China to the Caspian Sea with Tugay vegetation^11, 46^. Based on 20 different museum Caspian tiger specimens originating from Northwest China, Kazakhstan, Tajikistan, Uzbekistan, Turkmenistan, Azerbaijan, and Iran, only one major mtDNA haplotype was identified, which differed by only one nucleotide from Amur tigers across a 1.2 kb mitochondrial region; hence, synonymy between these two subspecies was proposed. However, in this study, analysis of the same samples used by Driscoll et al. (2009) at a genome-wide scale revealed close relatedness but distinctiveness of the Caspian and Amur tiger mitogenomes. In terms of nuclear genome, the Bengal tiger from the south contributed to approximately 26% of the nuclear genomic composition of the Caspian tiger via post-divergence gene flow, further suggesting a unique evolutionary origin and the subspecific distinctiveness of the Caspian tiger (VIR, *P. t. virgata*)

Three scenarios concerning the ancient colonization of Central Asia by tigers were previously postulated^11^: (i) a southern route via the Indian subcontinent, (ii) a northern route via the Siberian plain, and (iii) a historical “Silk Road” route through the Gansu corridor in Northwest China. Phylogenomic and demographic analyses and biogeographic modeling supported a possible initial expansion from East Asia to the modern range in Central Asia via the northern Siberian route (Route ii), followed by subsequent gene flow from the ancient Bengal tiger counterpart through the Himalayan corridor (Route i), previously proposed to be a biogeographical pass^10^.

Although the Caspian tiger shared a recent common ancestor with one South China tiger mitogenome, the lack of a continuous distribution of suitable habitat along the “Silk Road” from the LGM to the present excluded it (Route ii) as a potential a tiger dispersal pathway. The inclusion of distinctive mitochondrial lineages from the Amur, Caspian, and South China tigers in one clade (ALT/VIR/AMO, Fig. 2B) suggested a once-common ancestor of these mainland subspecies, whose ancient radiation center may have been located in East or Northeast Asia.

The taxonomy of the South China tiger is the most controversial among all, as the tiger is extinct in the wild and only survived by an inbred captive population derived from a small number of founders of uncertain origin. Early sampling of South China tigers uncovered two mtDNA lineages, one that is unique and basal to the other subspecies, and a second that is indistinguishable from Indochinese tigers^16, 20, 35, 47^. Here, for the first time, we retrieved population-level genetic information for 33 wild-origin South China tigers, from 30 of which mtDNA lineages were classified and 11 of which whole-genome data were obtained at 1-12× coverage, including the nominate type specimens. Nuclear genomic monophyly validated the South China tiger (AMO) as a distinct subspecies despite its mitochondrial paraphyly consisting of one unique basal haplogroup (AMO) and several others associated with the Malayan (JAX), Caspian (VIR), and Indochinese (COR) tigers. The nonrandom geographical distribution of mitochondrial haplogroups and genomic homogeneity across all South China tiger samples corroborated the “melting pot” hypothesis for eastern China, in which multiple historical populations met and merged. Modern South China tigers retained at least one relic, basal lineage once sheltered in Southwest China refugia, as projected by ENM, and several lines from the northbound populations expanding from Indochina.

Population genomic analyses of historical samples from the extinct tigers shed new light on tiger evolution from an ancient DNA perspective. Although D-statistics and admixture graph modeling revealed the occurrence of gene flow between tiger populations and mixed ancestry in some subspecies, genetic admixture mostly took place between populations with geographical proximity and most likely shortly after or during the early stage of divergence, while recent introgression was rare. The differentiation of modern tiger subspecies likely began with isolation driven by climate fluctuations during the Last Glacial Period c. 115,000 – c. 11,700 years ago, fortified through multiple range contraction–expansion–isolation cycles, and was completed after the last wave of post-LGM expansion.

China is the only country in the world that once harbored five out of the nine modern tiger subspecies or all those from mainland Asia, including the Amur (ALT) tiger in the northeast, the South China tiger (AMO) in the south and the east, the Indochinese tiger (COR) in the southwestern border adjacent to Southeast Asia, the Bengal tiger (TIG) along the southern fringe of Himalaya, and the Caspian tiger (VIR) in Northwest China. Ancient genomes showed that during the process leading to the phylogeographic partitioning and distinction of modern tiger subspecies, Southwest China may have served as a Late Pleistocene refugium for relic tiger lineage survival and eastern China as an ancient genetic melting pot for various lineages to meet and merge, as they expanded their range following the return of suitable habitats. The expansion of tigers across China en route to Northeast Asia and then westbound to Central Asia connected the northern subspecies such as the Amur and Caspian tigers with the southern mainland subspecies such as the Bengal and Indochinese tigers, highlighting the significance of mainland China as a stepping stone in the evolutionary history of tigers.

A thorough understanding of the tiger’s genomic diversity and evolutionary history is crucial for conservation and recovery planning for this charismatic megafauna. With genomic data retrieved from nowadays extinct tiger populations, this study resolved the basal rooting in the tiger phylogeny, illuminated cryptic biogeographic refugia and dispersal paths critical for the formation of extant landscape genetic patterns, and contributed to the much-needed scientific foundation guiding conservation.

## Materials and Methods

### Ancient specimens and radiocarbon dating

Twenty-five ancient felid skeletal samples were collected during paleontological excavations by the Far Eastern Branch of the Russian Academy of Sciences from limestone caves in the Primorsky Territory and the Jewish Autonomous Region of the RFE: Yunyh Speleologov Cave (43°29’N, 132°26’E), Letuchaya Mysh Cave (42°59’N, 133°05’E), Tetukhinskaya Cave (44°35’N, 135°36’E), and Koridornaya Cave (48°00’N, 130°59’E) (Tables 1, S2, Fig. S1). These specimens were identified to be from big cats based on morphological evaluation. Seven samples yielding genome sequencing data were selected for accelerator mass spectrometry (AMS)-^14^C radiocarbon dating (Table S1) at Peking University and Beta Analytic Co.. Calibration was performed based on the IntCal13 atmospheric curve^48^ with OxCal v4.2.4^49, 50^.

Historical samples, such as bones, pelts, and teeth, were assembled from 31 South China tigers located in various museums, institutes, or private collections in China, Europe, and USA (Table 1, S2, Fig. S1). All specimens were verified as having geographic origins in the traditionally recognized *P. t. amoyensis* (AMO) range, covering ten provinces, namely, Hubei, Hunan, Shaanxi, Yunnan, Guangdong, Jiangxi, Fujian, Shanxi, Sichuan, and Guizhou. The five nomenclatural type specimens of *P. t. amoyensis* (type locality: Hankou, China)^34^, including one holotype (museum ID: 3311) and four paratypes (3305, 3306, 3307, 3308) were sampled from Musée Zoologique de la Ville de Strasbourg, France, and incorporated in the dataset.

Six Caspian tiger specimens (VIR) were chosen from a previous museum collection^11^ for whole-genome sequencing based on mtDNA haplotype information reported in the study and represented the key *P. t. virgata* distribution in Uzbekistan, Kazakhstan, Turkmenistan, Tajikistan, and Azerbaijan (Tables 1, S2, Fig. S1).

### DNA extraction, library preparation, and targeted DNA capture

We processed degraded specimens in dedicated ancient DNA facilities at the School of Life Sciences, Peking University, China, and the Globe Institute, University of Copenhagen, Denmark. For DNA extraction, we evaluated and chose several optimized ancient DNA protocols based on sample type. Bone samples were digested in proteinase K buffer as previously described^51^. Tissue samples were digested with Buffer ATL from the DNeasy Blood & Tissue Kit (Qiagen, Valencia, CA, USA) with proteinase K. Pre-digestion with a small volume of digestion buffer at 37°C was applied to remove external DNA contamination^51, 52^, followed by a phenol‒ chloroform extraction protocol. Digest was then applied to a MinElute spin column (Qiagen) using a 10-fold volume of binding buffer^52^ and further processed following the manufacturer’s protocol.

Next-generation sequencing libraries were prepared with different methods optimized for ancient DNA. For DNA from relatively fresh samples, libraries were constructed with the NEBNext® Ultra™ II DNA Library Prep Kit for Illumina® (New England Biolabs, Ipswich, USA). DNA libraries for museum or archeological samples were built using a Blunt-End Single-Tube (BEST) method as previously described^53^. Libraries were sequenced on an Illumina HiSeq 2500 or X Ten platform at BIOPIC, Peking University, or Novogene Co., China. A ‘screen and boost’ sequencing strategy was used, in which 3-5 Gb of data from each library was retrieved, followed by subsequent boost sequencing according to the endogenous DNA content from the first round. For ancient RFE and Caspian tiger specimens, a Uracil-Specific Excision Reagent (USER, a mixture of uracil DNA glycosylase and the DNA glycosylase-lyase Endonuclease VIII) treatment was applied to the same batch of DNA used for library construction during the second round of boost sequencing.

For ancient RFE tiger samples, six samples with an endogenous DNA content above 0.5% were chosen for mitogenome capture enrichment. We used the tiger ‘myBaits Exper Mito’ kit (Daicel Arbor Bioscience, Ann Arbor, USA) following the manufacturer’s protocol v4.01 with the hybridization temperature set at 60°C. The DNA used for mitogenome capture was pre-treated with USER enzyme mix.

PCR amplification and Sanger sequencing of mitochondrial DNA fragments were performed for all historical South China tiger specimens following previously described procedures^12^. Partial mtDNA haplotypes and subspecies-diagnostic sites were compared to tiger voucher specimens^12, 16, 20^ to infer the maternal genetic ancestry of the individuals and to screen for samples suitable for further whole-genome resequencing.

## Data processing

### Quality control and alignment

Two separate data processing pipelines were implemented for the relatively recent museum specimens (∼100 years) of modern tiger subspecies (the Caspian and South China tigers) and the ancient RFE tiger specimens (>1,000 years). We used Cutadapter v2.1^54^ to trim sequencing adapters. Then 2 bp for modern samples or 5 bp for ancient samples were removed from both ends of the reads. Bases with a minimum quality of 30 at both ends and a minimum read length of 25 bp were kept for subsequent mapping. The retained reads were mapped to the tiger reference genome^13^. For modern specimens, we used the BWA-MEM^55^ algorithm to map the data. For ancient specimens, paired-end reads were collapsed with AdapterRemoval v2.2.2^56^, allowing a minimum length of 30 bp, and a BWA-backtrack algorithm was used for mapping with seeding disabled. The mapped paired-end reads were treated as two separate single-end reads during the removal of PCR duplicates. SAMtools v1.9^57^ was used to retain reads with a minimum mapping quality of 30 and remove PCR duplicates.

### Authentication of ancient DNA data

DNA damage patterns characteristic of ancient DNA were examined using mapDamage2^58^ for the RFE archeological specimens. All seven specimens with high-throughput sequencing data showed increased C to T and G to A substitutions at the 5’ and 3’ read ends. Pre- and post-USER mix treatment data for RUSA21 were compared (Fig. S2), and near complete removal of the damage pattern was observed suggesting the presence of ancient DNA. For ancient Russian tiger specimens, only libraries with USER mix treatment were included in the final dataset.

### Genotype calling

Single-nucleotide polymorphisms (SNPs) were identified using GATK v3.8 HaplotypeCaller and GATK v4.0 GenotypeGVCFs^59–61^ based on our dataset including 32 tigers from a previous study^20^. A hard filtering procedure was performed using BCFtools v1.9^62^ and homemade scripts based on the following criteria: (1) only biallelic SNPs were retained and SNPs within 5 bp of an indel were excluded; (2) BaseQRankSum (-1.96< BQRS < 1.96) and MappingQualityRankSum (MQRS > - 1.96) filters were applied; (3) genotype GT was marked as missing for loci in samples with a sequencing depth (DP) more than 4-fold of the average sample sequencing depth; (4) for rare alleles found only in degraded specimens, a minimum allele count of three and presence in at least three different specimens were required; (5) SNPs with a missing rate above 50% were excluded; and (6) only SNPs in scaffolds with a minimum length of 10 kb were included. The hard-filtered dataset was further filtered according to the needs of various downstream analyses and after considering variation in ancient DNA damage and sequencing depth among specimens.

### Autosomal neutral variant identification

We assembled autosomal regions by identifying and excluding the putative scaffolds related to the X and Y chromosomes (Fig. S7). Sex chromosome-derived scaffolds were identified by comparing the sequencing depth ratio between female and male tigers and the homologous scaffolds with the cat (*Felis catus*) sex chromosomes. In the first approach, only scaffolds with a minimum length of 10 kb were assessed. We identified scaffolds with a relative average sequencing depth in female individuals more than 1.5-fold of that in males as X chromosome derived and those less than 0.2× that in males as from the Y chromosome. Alternatively, we first applied a sliding window method to generate reads from the tiger reference genome^13^ with a window size of 100 bp and a step size of 50 bp. The reads from each scaffold were then mapped to the domestic cat sex chromosomes (Felis_catus_9.0, RefSeq assembly accession: GCF_000181335.3) using BWA v0.7.12 and a scaffold with mapping rate over 10% was considered sex-chromosome related. The results from the two methods were combined to obtain a total length of 111,848,290 bp (130,557,009 bp in *F. catus*) of X chromosome-related scaffolds and 1,481,767 bp (1,855,781 bp in *F. catus*) of Y chromosome-related scaffolds. The neutral autosomal regions were gathered after filtering out the coding regions and their flanking 1 kb from the recognized autosomal scaffolds.

### Mitochondrial DNA phylogeny

Mitochondrial DNA sequences were assembled independently with caution taken to exclude nuclear mitochondrial sequence (*numt*) contaminants^63, 64^. After passing quality control as described in the previous section, the raw reads as well as those from mitogenome capture enrichment were mapped to a previously confirmed cytoplasmic, *numt-*free mitogenome reference (NCBI ID: KP202268). The mapped reads were then processed in Geneious v9.1.5^65^ as input for iterative de novo assembly to further exclude *numt* contaminants. For the only sample with a relatively low sequencing depth (RUSA23, mitogenome coverage lower than 10×), we manually inspected and corrected the consensus sequence for *numt* contamination.

The highly variable control region was excluded from the analysis due to its unstable assembly results with too many ambiguous sites. We obtained three different mitogenomes from ancient RFE tigers, ten from South China tigers, and two from Caspian tigers.

We employed four different methods, as previously described^20^, to reconstruct the tiger mitogenome phylogeny, including neighbor joining (NJ), maximum parsimony (MP), maximum likelihood (ML) using PAUP v4.0b10^66^, and Bayesian methods using MrBayes v3.2.6^67^. Two *Panthera* species, the lion (*P. leo*, NCBI: KP202262.1) and snow leopard (*P. uncia*, NCBI: KP202269.1), and the clouded leopard (*Neofelis nebulosi*, NCBI: KP202291.1) the sister taxa of *Panthera*, were used as outgroups. TIM3+G+I was chosen as the optimal nucleotide substitution model by jModelTest v2.1.4^68^. The statistical support for the NJ, MP, and ML tree topologies was assessed in PAUP based on 1,000 bootstraps. Bayesian inference was performed with two independent Markov chain Monte Carlo (MCMC) runs, each set for 10,000,000 generations. Tree sampling was carried out every 1,000 generations and the first 25 percent of the iterations were discarded as burn-in.

### Whole-genome phylogeny

The whole-genome autosomal phylogeny was inferred based on transversion variants in the neutral regions, including the SNPs present only in outgroups. The dataset consisted of 3,838,774 sites that were concatenated into one consensus sequence for each sample. The phylogeny was then reconstructed with RAxML v8.2.11^69^ using the GTAGAMMA model and the node support was evaluated with 100 bootstrap replicates.

### Population genetic structure analysis

We performed PCA with *smartpca* in EIGENSOFT v6.1.4^70^ and calculated PCs using all the tiger samples with an average sequencing depth higher than 1×. The analysis was restricted to transversion sites in autosomal neutral regions to minimize the effect of ancient DNA damage. The final dataset consisted of 1,242,142 SNPs. The ancient RFE tiger specimen RUSA21 and other low-coverage South China tiger specimens were then projected with the *lsqproject* method implemented in *smartpca*.

We inferred population genetic structure with ADMIXTURE v1.3.0^71^ based on the same SNP panel used in PCA. The number of assumed ancestral populations (*K*) was set from 2 to 10. For each *K*, we ran 10 independent replicates with different starting seeds and chose the replicate with the highest likelihood.

### TreeMix phylogeny of population relationships

We used TreeMix v1.13^72^ to estimate the historical relationships among populations with admixture and infer possible migration events based on allele frequency in each population. The same panel of autosomal variants used for genome-wide phylogeny reconstruction was applied. To account for the effect of linkage disequilibrium, we set the *-k* flag at 1,000 to group nearby SNPs into blocks of 1,000 consecutive SNPs. Two historical South China tiger specimens (NGHE0002 and YANG0014), which displayed admixed genetic ancestry based on ADMIXTURE analysis, were designated as an independent population, “AMO_MIX”.

TreeMix runs were performed with the migration bands (*-m*) set from 0 to 7. Each migration band was run with 100 independent replicates using different starting seeds, and the replicate with the highest likelihood was presented.

### *D*-statistics estimation of gene flow

We applied *D*-statistics to infer possible gene flow between different tiger populations, specifically testing for excessive allele sharing as implemented in *qpDstat* of ADMIXTOOLS v5.1^73^. The analysis was conducted based on the same dataset used for whole-genome phylogeny reconstruction and the following scenarios were evaluated: (1) post-divergence gene flow in the Sumatran tiger (SUM) with all possible combinations of *D*(Outgroup, SUM, population 1 (pop1), population 2 (pop2)) calculated; (2) post-divergence gene flow in the joint population of the Indochinese tiger (COR) and Malayan tiger (JAX) with *D*(Outgroup, COR/JAX, pop1, pop2), and if any other population shared more alleles with these two tiger subspecies, then *D*(Outgroup, pop1, COR, JAX) was set; (3) post-divergence gene flow in the Bengal tiger (TIG) with *D*(Outgroup, TIG, pop1, pop2); and (4) post-divergence gene flow in the ancient RFE tiger (RUSA) with *D*(Outgroup, RUSA, pop1, pop2).

### Admixture graph modeling with *qpGraph*

We built admixture graphs using *qpGraph*^73^, which is based on *f*-statistics and takes into account genetic drift and mixture proportions, to model the evolutionary pathways and relationships among tiger populations. The same dataset including both modern and ancient tigers in D-statistics was included. We picked one sample with the highest sequencing depth to represent each population and searched the graph space using a heuristic algorithm implemented in *qpBrute*^74, 75^. A base graph was first constructed with two outgroups and the Sumatran tiger, followed by addition of the remaining populations in two rounds. The first round included three modern subspecies (JAX, TIG, and ALT) and one ancient population (RUSA) and resulted in 6,727 uniquely fitted graphs. The top three graphs with the lowest z-scores, the highest likelihoods, and no zero-length branches were set as the starting graph in the second batch, to which the remaining three subspecies (COR, AMO, and VIR) were added. Only one graph from the 15,179 unique fits without outliers or zero-length branches was presented as the best-fitting admixture graph model.

### Demographic inference with PSMC

We applied the PSMC^76^ model to infer the historical effective population size dynamics of various tiger subspecies, based on the same dataset used to calculate D-statistics. A consensus sequence was generated from the individual with the highest sequencing depth to represent each modern or ancient tiger population. Only the autosomal neutral region was included, and the rest of the genome was masked. The parameters for PSMC modeling were set as *-N25 -t20 -p “4+25*2+4+6”* and 100 bootstrap replicates were run to assess the robustness of the results. A mutation rate of 6.4×10^-9^ substitutions per site per generation was recalculated following the procedure described previously^20^, based on the autosomal neutral SNPs from this study, a generation time of five years for the tiger, and the overall divergence between the tiger and other *Panthera* species such as the lion and snow leopard.

### Demographic inference with G-PhoCS

The demographic history of tigers, including the historical effective population sizes, divergence times, and gene flow scenarios, was modeled in G-PhoCS^77^ based on the phylogeny inferred by TreeMix (Fig. S3 and Fig. S8). Two individuals with the highest sequencing depth were chosen to represent each modern tiger subspecies and one sample was from the ancient RFE population. The same dataset used for PSMC modeling was applied, and 44,092 autosomal loci were identified, each spanning l kb with a minimum inter-locus gap of 50 kb and a maximum missing rate of 5%. Four alternative combinations of individual and different outgroup species were evaluated to ensure model convergence. We first ran the model with all possible migration bands between different branches using 5,000 randomly chosen loci in two independent replicates. All migration bands were split into six groups to reduce the total number of parameters in each model. The total migration rate for each migration band was calculated as described in the G-PhoCS manual and migration bands with a total migration rate above 0.1 in at least one running batch were kept as the final model. We then ran the final model (Fig. S8) in two parallel replicates for each scenario of individual combinations. Estimates of model parameters (Fig. S9) were cross-checked for various runs and combined to obtained the final results. The divergence times for *P. tigris* and *P. leo* at 3.72 Mya (95% CI: 2.04–7.60 Mya) and *P. tigris* and *P. uncia* at 2.67 Mya (95% CI: 1.14-5.92 Mya) were used as calibration values for the model^2^. A generation time of five years for the tiger was applied.

### Mitogenome coalescent time estimation

The coalescent time of modern and ancient tiger mitogenomes was estimated in BEAST v2.5.2^78^ based on the same nucleotide substitution model, TIM3+I+G, used in phylogeny inference, a strict-clock setting, and a coalescent–exponential– population model. The coalescent times between *N. nebulosa* and *Panthera* spp. (6.37 Mya, 95% CI: 4.47-9.32 Mya) and between *P. leo* and *P. uncia*/*P. tigris* (3.72 Mya, 95% CI: 2.04-7.60 Mya) were used as calibration values. MCMC was performed with five independent runs for 50,000,000 iterations each. Sampling was performed every 1,000 generations and the first 20% of values were discarded as burn-in.

### Ecological niche modeling

We built ecological niche models in Maxent v3.4.0^79^ to correlate presence data with the species’ ecological requirements and to project tiger habitat suitability during different geological periods. A total of 781 tiger locality points (Fig. S5) were assembled from published studies^10^ and records from the IUCN Cat Specialist Group^80, 81^. Bioclimate variables from WorldClim 1.4^82^ were used as predictors and eight of the 19 variables (Table S4) were retained after removing the highly correlated ones (|”| ≥ 0.8). The models were built at a resolution of 10 arc-minutes. The spatial extent ranged from 20° W to 144° E and 10° S to 82° N and was restricted to the region with tiger occurrence in order to reduce biased model fitting. The models were constructed under the modern climate with an area under the curve (AUC) of 0.835 and then projected to three types of, namely, the LIG period (c. 120,000-140,000 BP), the LGM (c. 22,000 BP), and the Mid-Holocene (c. 6,000 BP).

Two additional suitable habitat layers for tigers were generated based upon the published biome and vegetation layers for the modern^83^ and LGM^84^ geological periods. The binary suitability (suitability vs. unsuitability) of different biomes and vegetation types for tigers (Table S4) was manually designated according to tigers’ preferred habitat classification as listed on the IUCN Red List of Threatened Species^17^.

## Acknowledgments

All samples were recruited in compliance with the Convention on International Trade in Endangered Species of Wild Fauna and Flora (CITES) through permissions issued to the School of Life Sciences (PI: S.-J.L.), Peking University, by the State Forestry Administration of China. We thank all the collaborators, institutes, and zoos that provided the specimens listed in Table S2 upon which this study is based. Special thanks are given to the following people who provided important help during various stages of the project: Zhou Xiaobo, Liao Lijing, Wu Jiayan, Feng Chengli, Xiang Shuanglin, <u>Shen Youhui</u>, Xie Can, Zhang Li, Chen Youling, Tang Fuchou, Enrico Cappellini, Meaghan Mackie, Miao Lin, Hu Xuesong, Huang Ji, Yu He, Meng Hao, Fu Qiaomei, Eleanor Hoeger, Marisa Surovy, Neil Duncan, Sara Ketelsen, Marie-Dominique Wandhammer, Virginia Rakotondrahaja, Alexei Abramov, Igor Ya. Pavlinov, Elena I. Zholnerovskaya, Natasha V. Lopatina, Gu Xiaodong, Gu Haijun, Dale Miquelle, and David Smith. We would also like to pay tribute to the late Ulysses Seal and Peter Jackson for their dedication in tiger conservation and pioneer effort in assembling voucher specimens for genetic study. This work was supported by the National Natural Science Foundation of China (NSFC 32070598) and the Peking-Tsinghua Center for Life Sciences. M.P.T. was funded, in part, by the Russian Foundation for Basic Research (project 18–04–00327).

## Data Availability

The Next Generation Sequencing raw data of the tiger samples have been deposited in the Sequence Read Archive (BioProject ID: PRJNA822019).

## Notes

### Competing Interest Statement

The authors have declared no competing interest.

