## Supplemental figures for "Ancient DNA Reveals China as a Historical Genetic Melting Pot in Tiger Evolution"

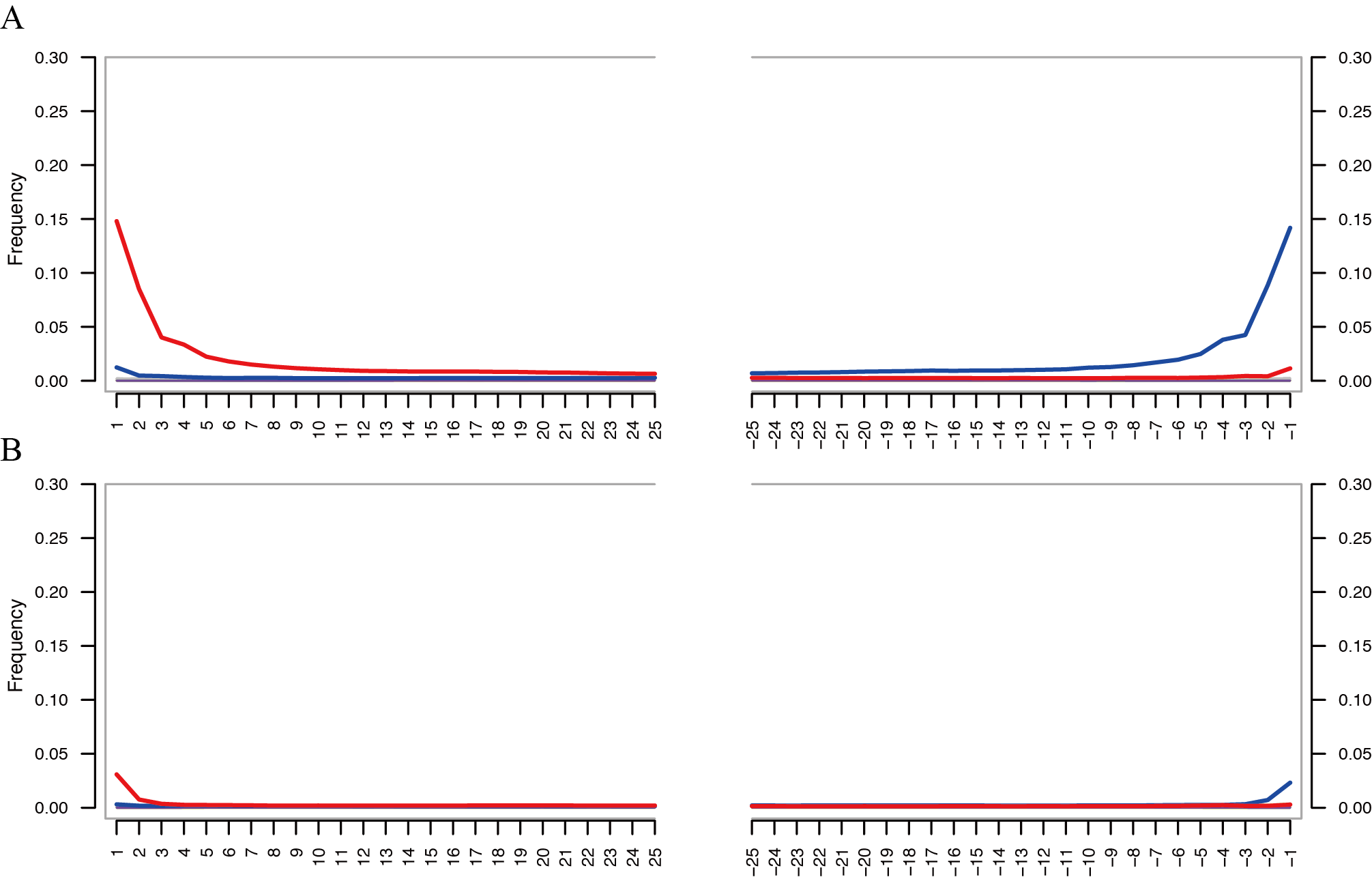

**Fig. S1. Authentication of ancient DNA sequencing data from RUSA0021.** Panels A and B show the different DNA substitution patterns at the 5' (left) and 3' (right) ends of reads from RUSA0021 before and after USER mix treatment, respectively. Red lines refer to C-to-T substitutions, and blue lines refer to G-to-A substitutions.

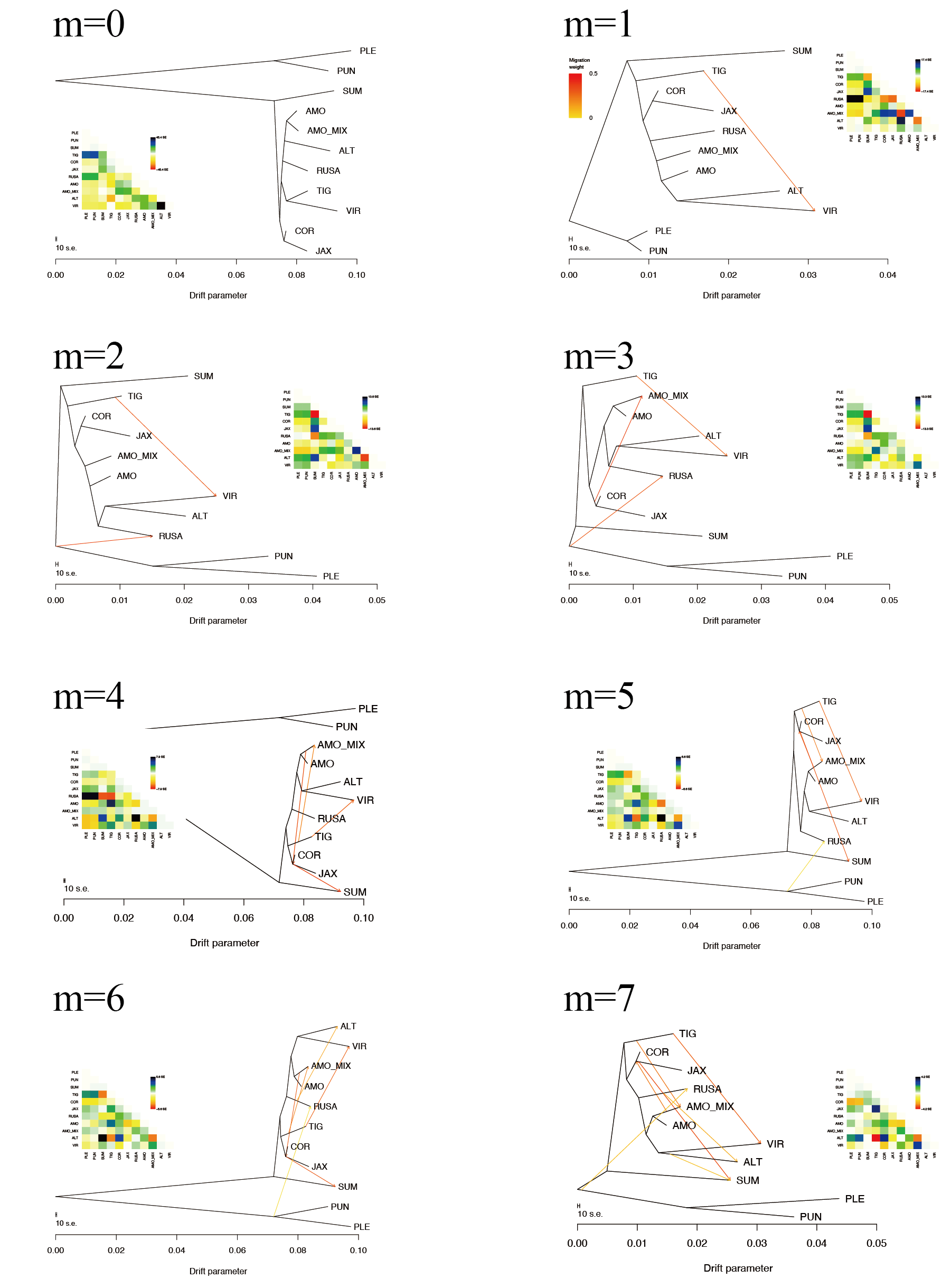

**Fig. S2. TreeMix phylogeny of ancient and modern tigers.** Samples were grouped by modern subspecies or ancient RFE population. South China tigers with admixed ancestry were assigned as a separate group (AMO_MIX). Migration edges (m) were inferred from 0 to 7. The migration band inference results were similar to the results for gene flow and population admixture obtained with *D*-statistics and admixture graph modeling. We used the topology with 7 inferred migration bands for further demographic history modeling.

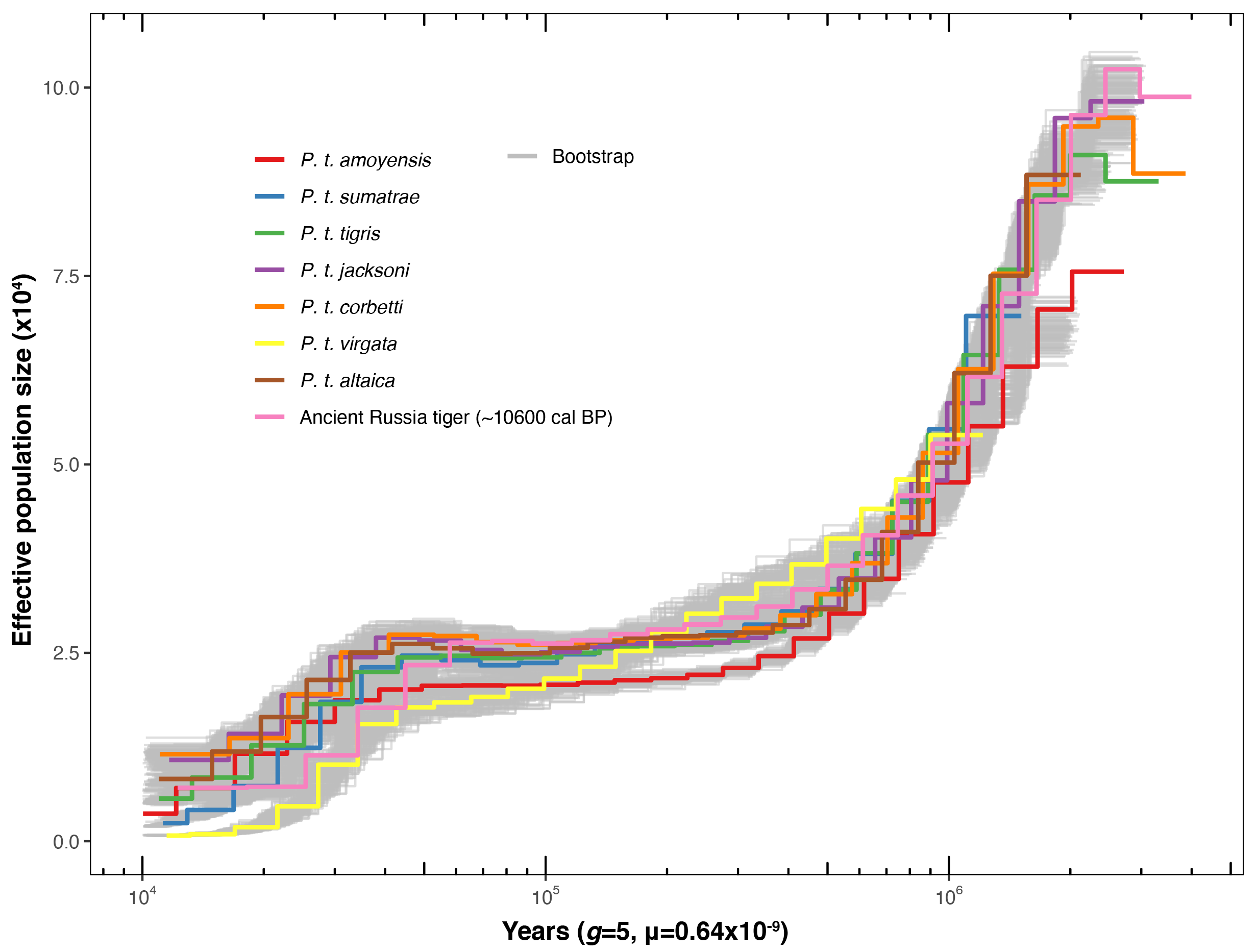

**Fig. S3. Demographic history analysis for different tiger subspecies estimated in PSMC.** PSMC was applied for each tiger, and one for each subspecies is shown here. The generation time *g* was set to 5 years, and the mutation rate $\mu$ was calculated to be 0.64$\times$10^-9^ substitutions per site per year. Support values from 100 bootstrap replicates for each run are shown in gray.

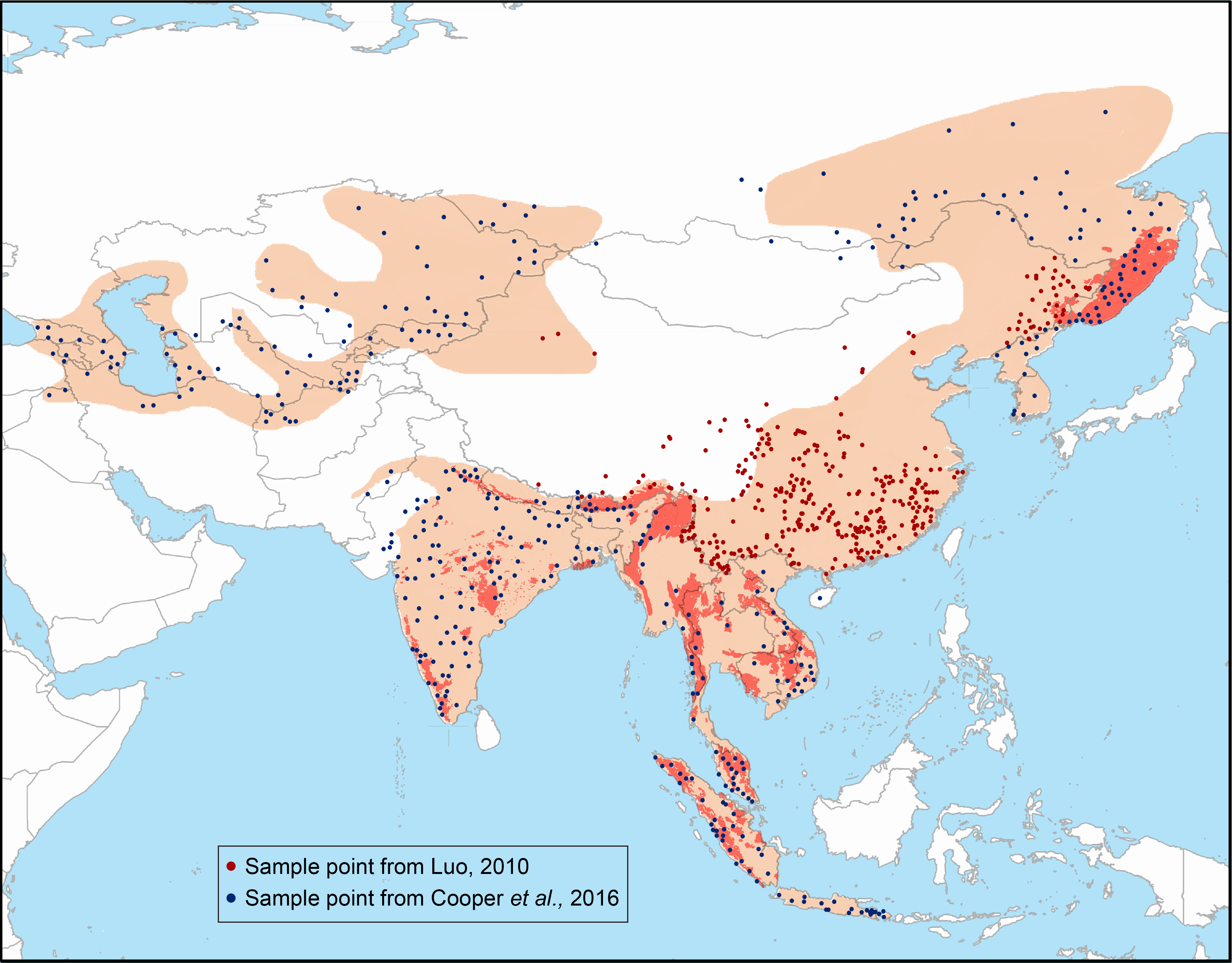

**Fig. S4. Tiger locality data in modern times are color-coded by data source and the range distribution background used for ecological niche modeling (ENM) in Maxent.** Blue dots are tiger presence localities used in a previous study^10^, and red dots are data assembled from the literature^80,81^ in this study with a special focus on the distribution in China to fill the data gap in previous studies.

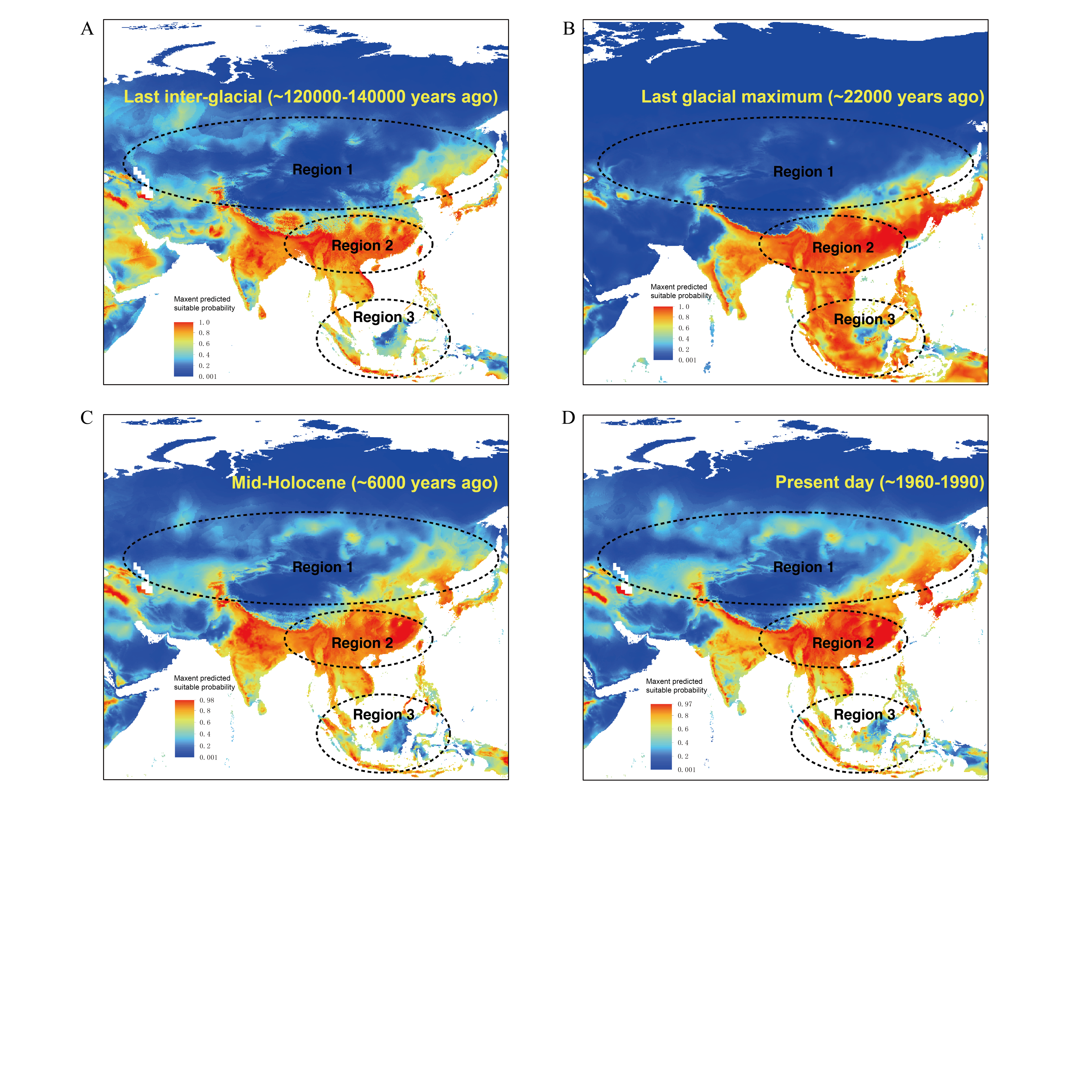

**Fig. S5. Ecological niche modeling results of the tiger distribution during four different periods.** (*A*) The Last Interglacial period (LIG, approximately 120,000-140,000 years ago), (*B*) the Last Glacial Maximum (LGM, approximately 22,000 years ago), (*C*) the mid-Holocene (approximately 6,000 years ago), and (*D*) present day (data from approximately 1960-1990).

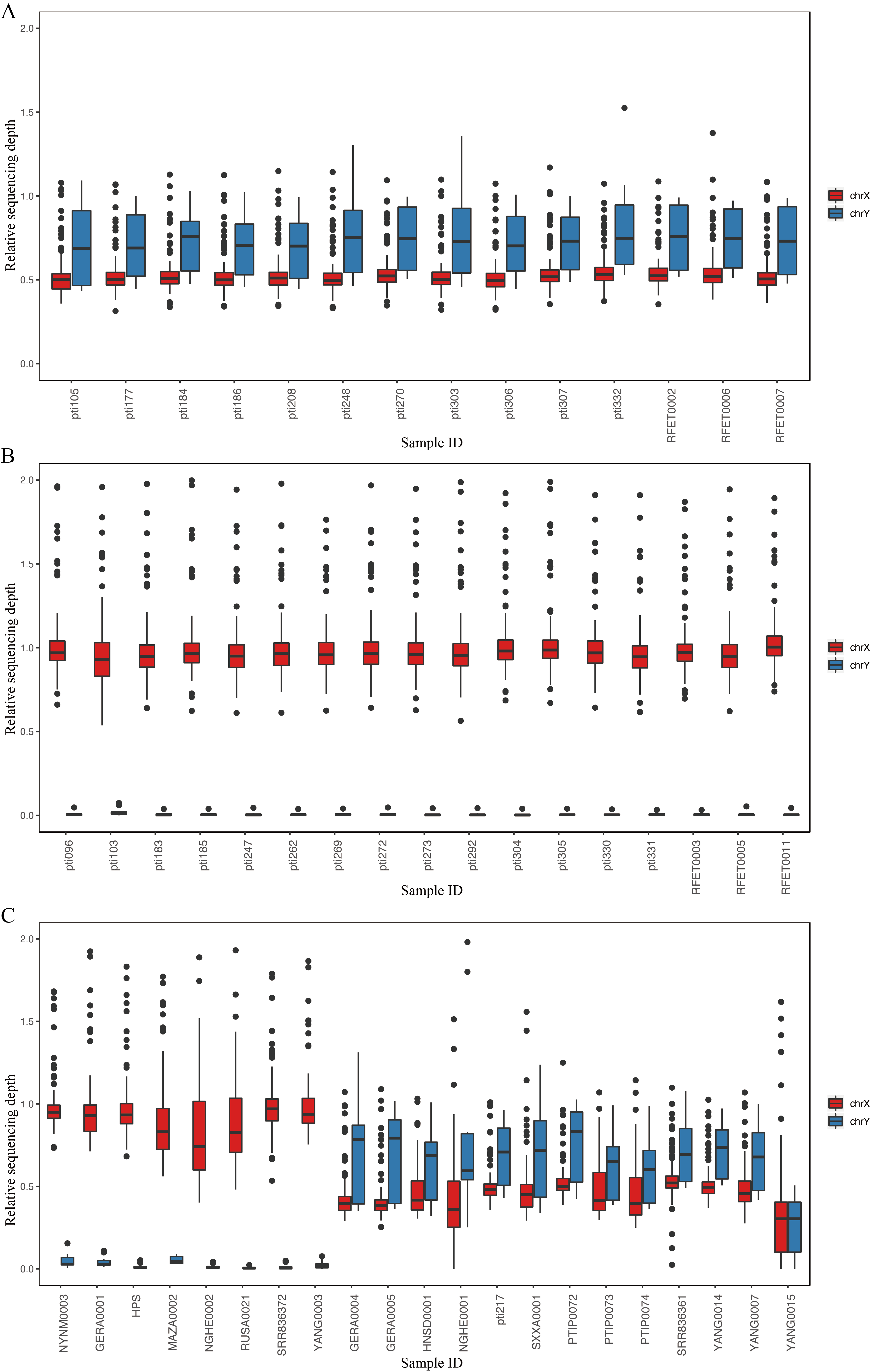

**Fig. S6. Detection of sex-chromosome-related scaffolds and sex inference for the samples.** Relative sequencing depth of sex chromosomes for male (*A*) and female (*B*) individuals in this study. (*C*) Sex inference based on the relative sequencing depth of sex chromosomes.

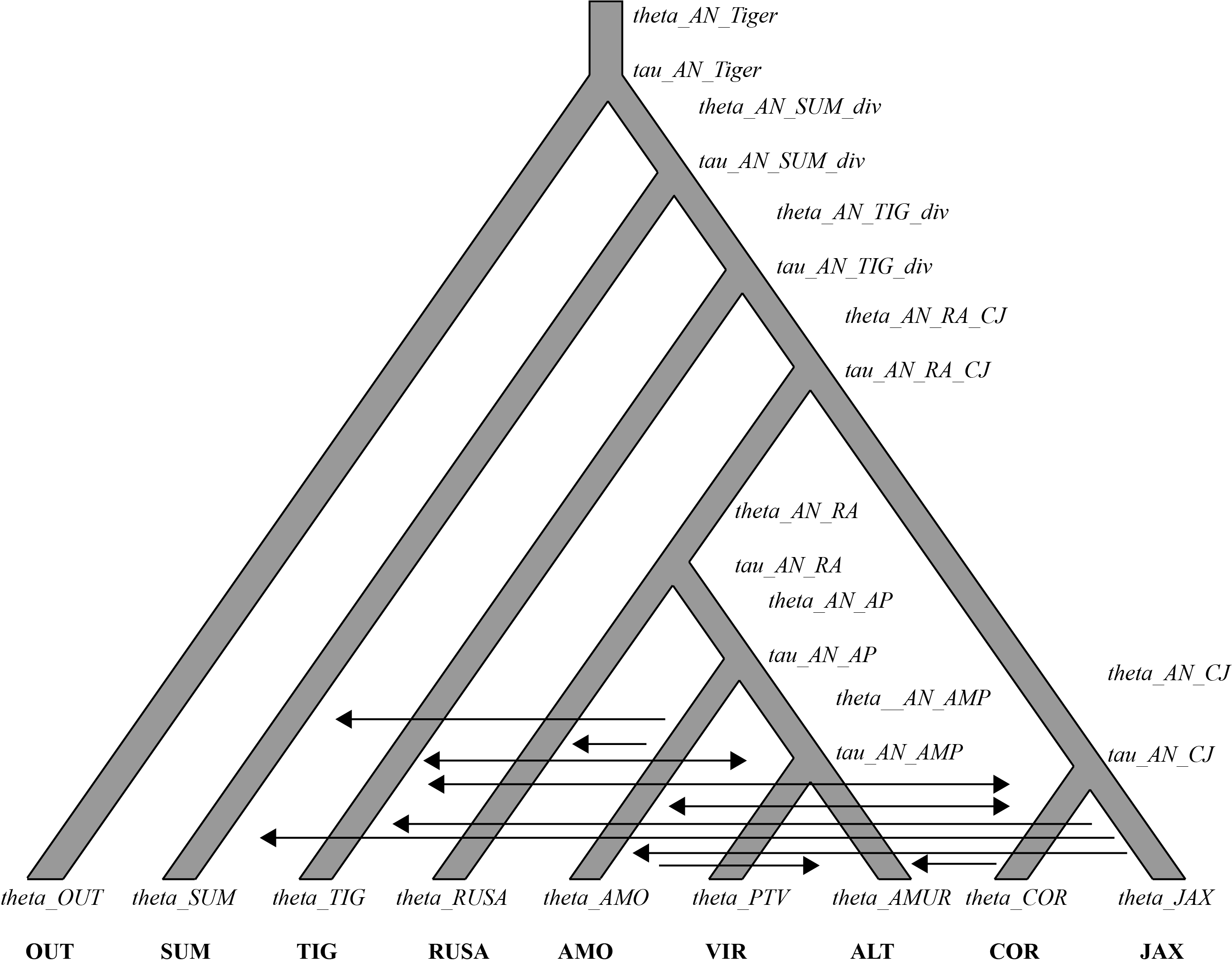

**Fig. S7. G-PhoCS modeling parameters and migration bands between different populations.** The parameter *theta* refers to the effective population size, and *tau* refers to the coalescence time of the populations from the tip to the node.

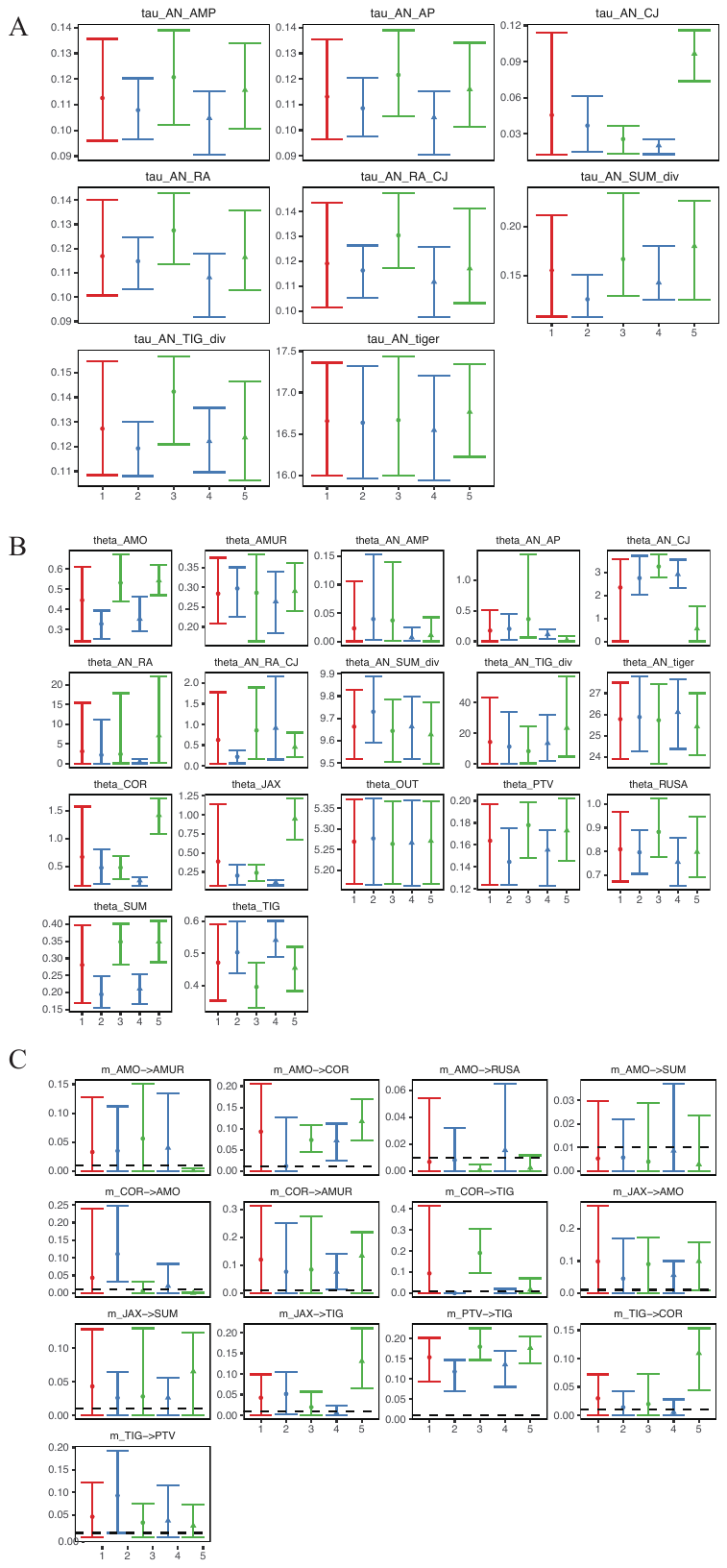

**Fig. S8. Parameter estimation result from G-PhoCS.** The numbers on the x-axis refer to different batches of replicates, except for 1, which indicates the summarized result from all replicates. Parameter estimates include (*A*) *tau*, (*B*) *theta,* and (*C*) the total migration rate *m*.

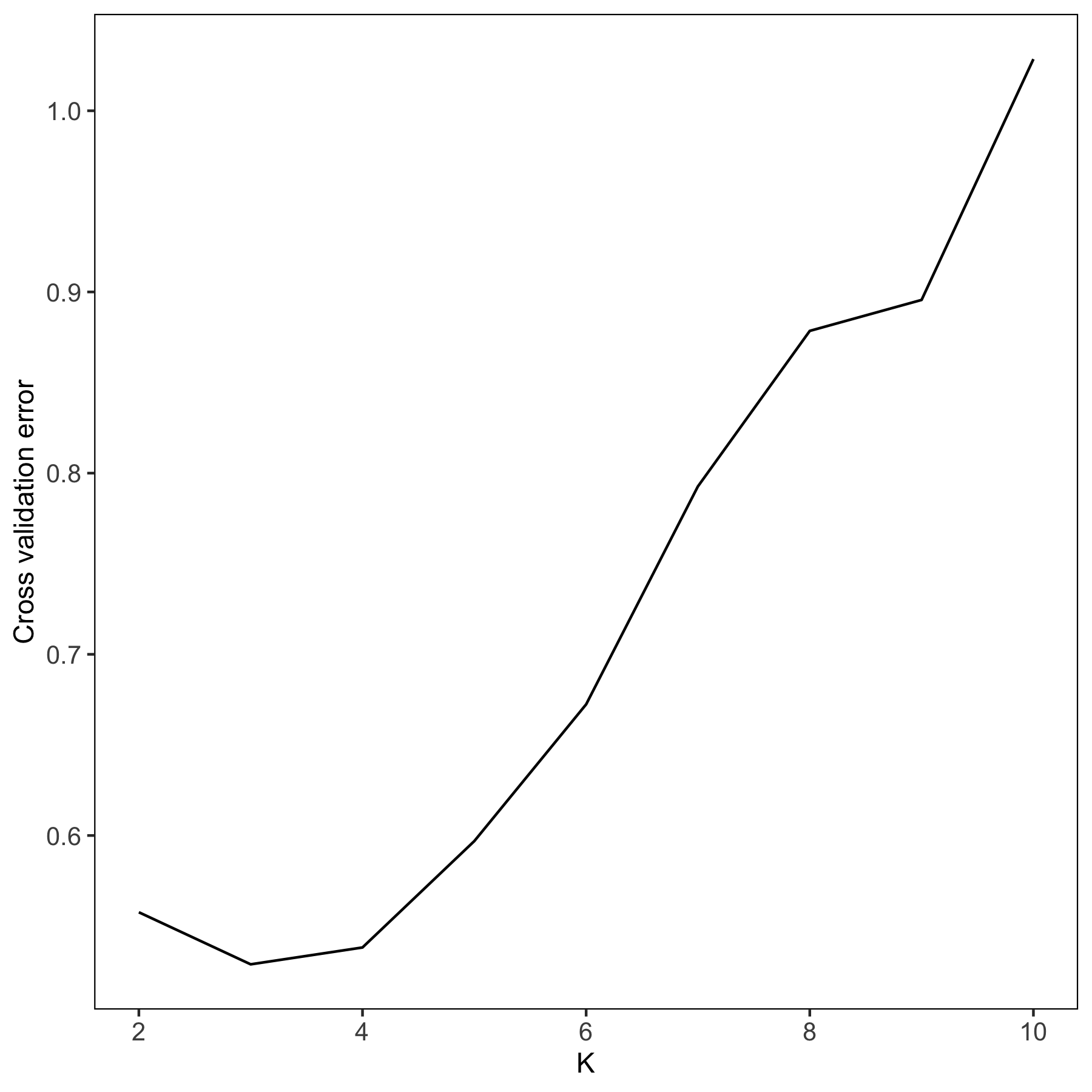

**Fig. S9. The cross-validation (CV) error for admixture runs with different numbers of ancestral populations assumed (*K)***.
